# Goblet Cell Hyperplasia Increases SARS-CoV-2 Infection in COPD

**DOI:** 10.1101/2020.11.11.379099

**Authors:** Jaspreet K. Osan, Sattya N. Talukdar, Friederike Feldmann, Beth Ann DeMontigny, Kailey Jerome, Kristina L. Bailey, Heinz Feldmann, Masfique Mehedi

**Affiliations:** Department of Biomedical Sciences, University of North Dakota School of Medicine & Health Sciences, Grand Forks, ND 58202, USA; Divison of Intramural Research, Rocky Mountain Laboratories, National Institute of Allergy and Infectious Diseases, National Institutes of Health, Hamilton, MT 59840, USA; Department of Internal Medicine, Pulmonary, Critical Care and Sleep and Allergy, University of Nebraska Medical Center, Omaha, NE 68198, USA

**Keywords:** SARS-CoV-2, COVID-19, goblet cells, ciliated cells, COPD, squamous metaplasia, air-liquid interface, syncytium, cell sloughing, goblet cell hyperplasia.

## Abstract

SARS-CoV-2 has become a major problem across the globe, with approximately 50 million cases and more than 1 million deaths and currently no approved treatment or vaccine. Chronic obstructive pulmonary disease (COPD) is one of the underlying conditions in adults of any age that place them at risk for developing severe illness associated with COVID-19. We established an airway epithelium model to study SARS-CoV-2 infection in healthy and COPD lung cells. We found that both the entry receptor ACE2 and the co-factor transmembrane protease TMPRSS2 are expressed at higher levels on nonciliated goblet cell, a novel target for SARS-CoV-2 infection. We observed that SARS-CoV-2 infected goblet cells and induced syncytium formation and cell sloughing. We also found that SARS-CoV-2 replication was increased in the COPD airway epithelium likely due to COPD associated goblet cell hyperplasia. Our results reveal goblet cells play a critical role in SARS-CoV-2 infection in the lung.

## Introduction

Severe acute respiratory syndrome coronavirus 2 (SARS-CoV-2, a causative agent of coronavirus disease 2019, COVID-19) that emerged in December 2019 in Wuhan, China. Since then, this pathogen has caused havoc in the healthcare systems worldwide and consequentially ravaged the economy of countries with COVID-19 outbreaks. There is currently no FDA-approved vaccine against SARS-CoV-2. SARS-CoV-2 is a nonsegmented, positive-sense, single-strand RNA virus that causes both upper and lower respiratory tract infections. Most patients exhibit fever and cough, and a subset of patients advance to severe acute respiratory distress syndrome (ARDS) (Guan et al., 2020; Yang et al., 2020). Therefore, patients with underlying chronic obstructive pulmonary disease (COPD) are vulnerable to COVID-19, and in fact, COPD is one of the high-risk factors for severe illness associated with COVID-19 (CDC, 2020; Leung et al., 2020; Sin, 2020).

Viral infections begin by the attachment of viral particles to entry receptors on the host cell. The tissue expression and distribution of the SARS-CoV-2 entry receptor angiotensin-converting enzyme 2 (ACE2) and its co-factor transmembrane serine protease 2 (TMPRSS2) determine the tropism of virus infection (Hoffmann et al., 2020; Li et al., 2003), and viral infection in human airway epithelium depends on ACE2 expression (Hamming et al., 2004; Jia et al., 2006). For successful entry into cells, SARS-CoV-2 uses the serine protease TMPRSS2 for S protein priming (Hoffmann et al., 2020). ACE2 is highly expressed in the small intestine, testis, kidneys, heart, thyroid, and adipose tissue and is expressed at moderate expression levels in the lung, colon, liver, bladder, and adrenal gland; and lowest in the blood, spleen, bone marrow, brain, blood vessels, and muscle (Hamming et al., 2004; Li et al., 2020). ACE2 expression in the lungs is predominantly observed in alveolar type 2 (AT2) cells (Lukassen et al., 2020; Qi et al., 2020; To and Lo, 2004; Ziegler et al., 2020), but ciliated cells also express ACE2 in the respiratory epithelium (Sims et al., 2005). Recent RNAseq-based studies have suggested that ACE2 is more highly expressed on goblet cells in the nasal airways and on secretory cells in subsegmental bronchial branches of the lung (Lukassen et al., 2020; Sungnak et al., 2020; Ziegler et al., 2020). Although ACE2 and TMPRSS2 expressions are higher in nonciliated goblet cells compared to ciliated cells (Lukassen et al., 2020; Sungnak et al., 2020; Zhang et al., 2020; Ziegler et al., 2020), it appears that goblet cells are underappreciated in the SARS-CoV-2 infection studies. The possibility that SARS-CoV-2 infects goblet cells could explain the presence of viral RNA in sputum (Wang et al., 2020) and might explain the efficient transmission of the virus from person to person (Dhand and Li, 2020; Wolfel et al., 2020). Importantly, goblet cell hyperplasia is a characteristic pathological feature of COPD patients, who are vulnerable to severe disease associated with COVID-19 (Lippi and Henry, 2020; Shimura et al., 1996; Zhao et al., 2020). Therefore, it is prudent to determine to what extent SARS-CoV-2 infects goblet cells in the lung.

To determine the expression of the SARS-CoV-2 receptor and its preferential cell tropism in the lung, we developed an in vitro airway epithelium model by differentiating primary normal human bronchial (NHBE) cells derived from either a patient with COPD or a healthy adult (non-COPD). The COPD airway epithelium model recapitulates many bronchial characteristics of COPD. We evaluated the expression of ACE2 and TMPRSS2 and studied SARS-CoV-2 infection in these in vitro airway epithelium models. We found that SARS-CoV-2 primarily infects nonciliated goblet cells due to high expression of both ACE2 and TMPRSS2 in these cells. Goblet cell hyperplasia increases of SARS-CoV-2 infection in the COPD airway epithelium. Thus, SARS-CoV-2 targeting and replication in goblet cells may explain the development of more severe COVID-19 in COPD patients.

## Results

### The airway epithelium model recapitulates the chronic bronchial characteristics of COPD

We first established an in vitro airway epithelium model by differentiating NHBE cells from either a healthy adult or a COPD patient (deidentified) at the air-liquid interface (ALI). We found that 4 weeks of differentiation provides a fully differentiated pseudostratified mucociliary airway epithelium for both conditions (Figures 1A and B), which allows the use of this model for side-by-side comparisons. NHBE cells from a healthy adult differentiated primarily into a pseudostratified columnar epithelium, whereas the NHBE cells from a COPD patient differentiated into a mix of pseudostratified and stratified columnar epithelium that contained all three main cell types of the respiratory epithelium (Pawlina, 2016; Rayner et al., 2019; Rigden et al., 2016): ciliated cells, goblet cells, and basal cells (Figure 1B). We found that the apical site of the epithelium mainly consists of ciliated and nonciliated goblet cells (Figures 1C and D). Studies have shown that mucin 5AC (MUC5AC) is predominately expressed by airway goblet cells and that mucin 5B (MUC5B) is expressed by goblet cells of submucosal glands (Whitsett, 2018). Club cells are the progenitors of goblet cells, which might express both MUC5AC and MUC5B (Kiyokawa and Morimoto, 2020; Okuda et al., 2019). As expected, we found heterogenicity in the cell population with differential expression of common goblet cell markers (Figure 1E) (Lukassen et al., 2020). Basal cells are known for their self-renewal property and give rise to multiple types of differentiated airway epithelial cells (Crystal, 2014). For the detection of basal cells, we sectioned the epithelium and stained the sections for the basal cell marker P63 (Persson et al., 2014; Wang et al., 2002). We found that basal cells reside at the basal membrane of both epithelia and that the COPD epithelium has more basal cells than the healthy epithelium as it is known from respiratory epithelium of COPD patients (Figure 1F) (Higham et al., 2019; Polosukhin et al., 2011).

**Figure 1.**
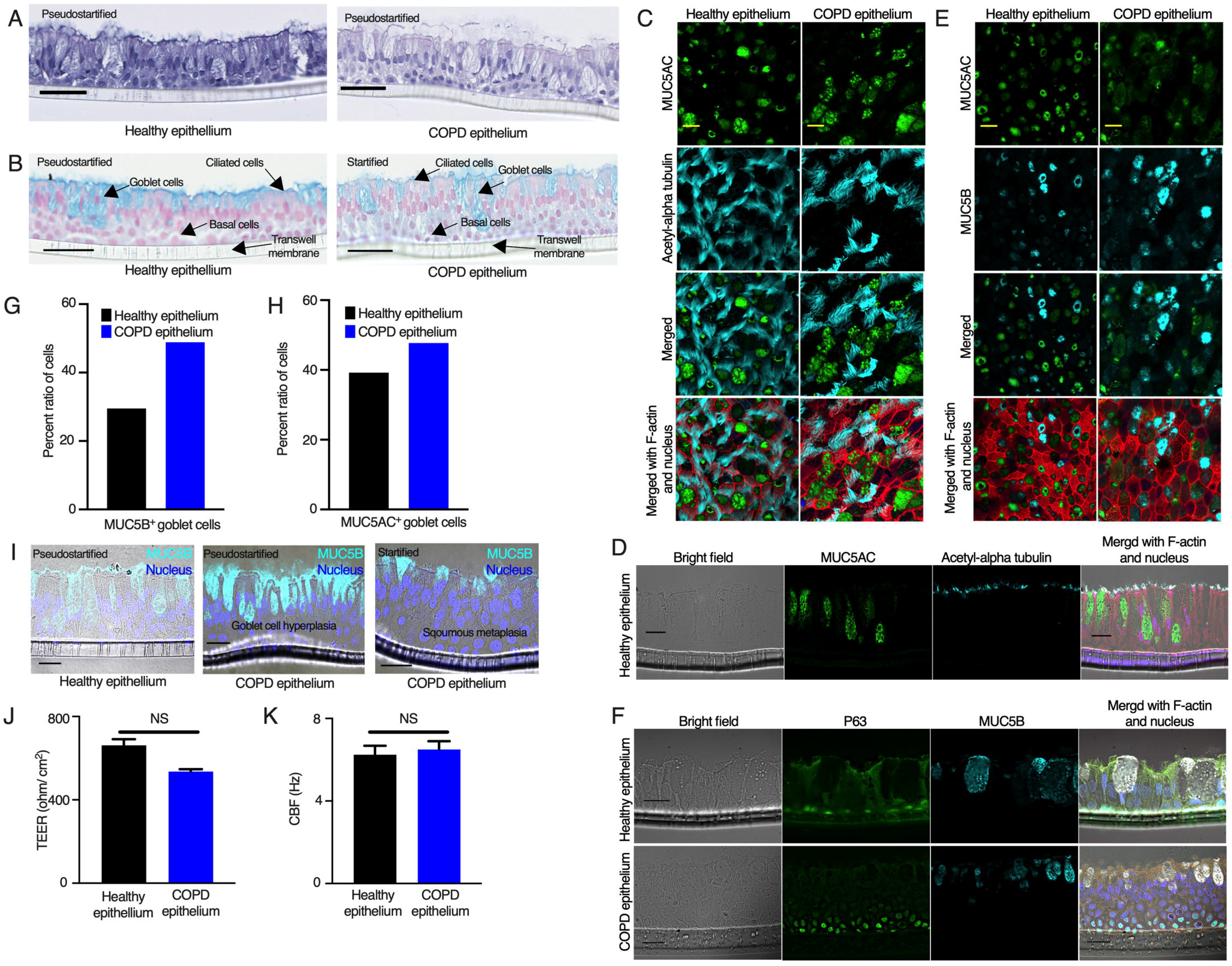
The airway epithelium model recapitulates the bronchial characteristics of COPD. **(A).** Airway epithelium derived from differentiated NHBE cells obtained from a healthy adult or COPD patient were fixed, embedded in paraffin, sectioned and stained with hematoxylin and eosin (H&E). The nuclei are stained dark-purple, and the cytoplasmic components are pink. Bar = 50 μm. **(B)**. The sectioned epithelia were stained with Alcian blue. Mucosubstances are stained blue, whereas the cytoplasmic components are pale pink, and the nuclei are pink/red. Bar = 50 μm. **(C).** The apical sites of the airway epithelia were fixed, permeabilized and stained for the goblet cell marker MUC5AC (anti-MUC5AC, green), the ciliated cell marker acetyl-alpha tubulin (anti-acetyl-alpha tubulin, cyan), and F-actin (rhodamine phalloidin, red); the nuclei were also stained with DAPI (blue). Bar = 10 μm. **(D).** The sectioned epithelia were stained for the ciliated cell marker acetyl-alpha tubulin (anti-acetyl-alpha tubulin, cyan), the goblet cell marker MUC5AC (anti-MUC5AC, green), and F-actin (rhodamine phalloidin, red); the nuclei were also stained with DAPI (blue). Bar = 20 μm. **(E).** The apical sites of the airway epithelia were fixed and stained for two goblet cell markers, MUC5AC and MUC5B [anti-MUC5AC (green) and anti-MUC5B (cyan), respectively] and F-actin (rhodamine phalloidin, red); the nuclei were also stained with DAPI (blue). Bar = 10 μm. **(F).** The sectioned epithelia were stained for the basal cell marker P63 (anti-P63, green), the goblet cell marker MUC5B (anti-MUC5B, cyan), and F-actin (rhodamine phalloidin, red); the nuclei were also stained with DAPI (blue). Bar = 20 μm. **(G).** The MUC5B^+^ goblet cells were counted from tiled images of the airway epithelium stained for MUC5B (anti-MUC5B, cyan) (described above) obtained with a Leica DMi8 epifluorescence microscope. A total of approximately 3,500 cells were counted from each epithelium to determine the ratio. **(H).** The MUC5AC^+^ goblet cells were counted from random fields in confocal images of the airway epithelium stained for MUC5AC (anti-MUCAC, cyan) (described above). A total of approximately 1,500 cells were counted from each epithelium to determine the ratio. **(I).** The sectioned epithelia were stained for MUC5B (anti-MUC5B, cyan); the nuclei were stained with DAPI (blue). Bar = 20 μm. **(J).** Transepithelial electrical measurements (TEERs) of the airway epithelia were obtained. The data were obtained by combining three independent Transwell reads, and each Transwell read was an average of three independent reads. The error bars represent the SEMs. **(K).** The ciliary beat frequency (CBF) was measured on the airway epithelia. The data were obtained by combining three independent Transwell reads, and each Transwell read was an average of six random point reads. The error bars represent the SEMs. The statistical significance was determined by unpaired two-tailed t-tests. The results from one independent experiment are shown.

COPD is associated with abnormal airway and alveolar responses during exposure to noxious stimuli (Brusselle et al., 2011). Because our COPD airway epithelium model was differentiated from NHBE cells, it should recapitulate the bronchial airway phenotypes instead of the alveolar phenotype as more commonly associated with emphysema (Barnes, 2013). To determine whether our in vitro COPD airway epithelium model recapitulates some of the bronchial pathophysiological characteristics of COPD, we focused on two different aspects: goblet cell hyperplasia and squamous metaplasia. First, we compared the number of MUC5AC^+^ or MUC5B^+^ goblet cells between healthy and COPD epithelia. We found a higher ratio of goblet cells in the COPD airway epithelium (Figures 1G and H). The higher number of goblet cells in the COPD epithelium suggests a persistent goblet cell differentiation, which results in goblet cell hyperplasia (Kim et al., 2015; Reid et al., 2018; Shaykhiev, 2019). Indeed, we found a patch of higher number of goblet cells with extensive mucus secretion in the COPD epithelium (Figure 1I, center). We also found an apparent loss of pseudostratified epithelium accompanied by squamous metaplasia in the COPD epithelium, which is a common pathological phenotype in COPD (Figure 1I, right) (Rigden et al., 2016).

To determine the biophysical properties of these respiratory epithelia, we assessed the tissue membrane integrity (transepithelial electrical resistance, TEER) and ciliary function (ciliary beat frequency, CBF) and found no significant difference in these biophysical properties between healthy and COPD epithelia (Figures 1J and K). These results indicated that NHBE cells from patients with COPD produced a mix of pseudostratified and stratified highly differentiated mucociliary epithelium with appropriate biophysical properties.

### ACE2 and TMPRSS2 are expressed at higher levels in goblet cells

SARS-CoV-2 infects the human airway epithelium, and virus entry depends on the host cell receptors ACE2 and its co-factor TMPRSS2 (Hoffmann et al., 2020). We quantified the ACE2 transcript levels in a lung epithelial cell line (A549 cells) and primary NHBE cells in a monolayer or in the differentiated airway epithelium by real-time PCR. We did not detect ACE2 transcripts in the A549 cells (Figure S1A), which might indicate low or no expression of ACE2 confirming previous data (Blanco-Melo et al., 2020; Harcourt et al., 2020; Hoffmann et al., 2020; Jia et al., 2005). However, we detected ACE2 transcripts in primary NHBE cells in both the monolayer and differentiated airway epithelium (Figure S1A). Despite their similar biophysical properties (tissue barrier integrity and ciliary function, Figures 1J and K, respectively), ACE2 expression was higher in the COPD epithelium compared to those derived from a healthy donor (Figures S1A-C). We then visualized the expression of ACE2 in the airway epithelium by immunofluorescence imaging. We observed ACE2 expression in both the healthy and COPD airway epithelia and found that ACE2 expression hardly overlapped with cilia on the apical site of the epithelium (Figures 2A and S1D). Nevertheless, low levels of ACE2 expression were observed on ciliated cells in our model (Figure S1E). These results suggest that the SARS-CoV-2 entry receptor ACE2 is mainly expressed on non-ciliated cells in the respiratory epithelium.

**Figure 2.**
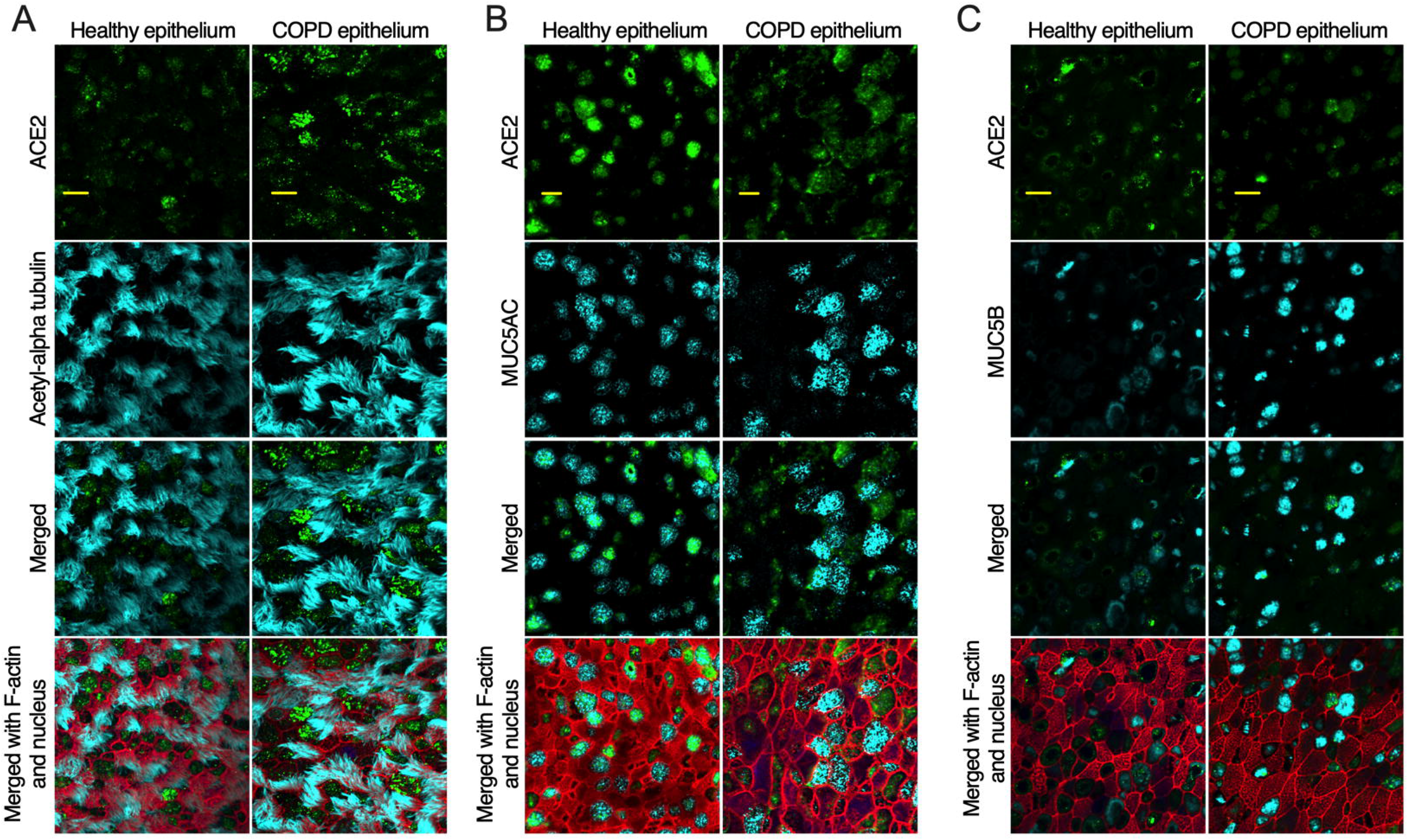
ACE2 is expressed at higher levels in goblet cells. The airway epithelia were generated and fixed as described in Figure 1. **(A)** Cells were stained for ACE2 (anti-ACE2, green), cilia (anti-acetyl-alpha tubulin, cyan) F-actin (rhodamine phalloidin, red) and nuclei (DAPI, blue). **(B).** Cells were stained for ACE2 (anti-ACE2, green), MUC5AC+ goblet cells (anti-MUC5AC, cyan) F-actin (rhodamine phalloidin, red) and nuclei (DAPI, blue). **(C).** Cells were stained for ACE2 (anti-ACE2, green), MUC5B+ goblet cells (anti-MUC5B, cyan), F-actin (rhodamine phalloidin, red) and nuclei (DAPI, blue). Bar =10 μm. See also Figures S1 and S2.

Because ACE2 staining hardly overlapped with acetyl-alpha-tubulin, we tested the expression of ACE2 along with that of the goblet and club cell markers MUC5AC and MUC5B (Lukassen et al., 2020). Indeed, ACE2 overlapped with MUC5AC (Figure 2B) and MUC5B (Figures 2C and S1F). We also compared ACE2 expression with the expression of E-cadherin and Zonula occludens-1 (ZO-1), markers for adherens junction and tight junction proteins, respectively. The results showed that ACE2 expression did not overlap with apical tight junctions or adherent junctions (Figure S2), which suggests that ACE2 expression is primarily located within the cellular boundary and does not impact the tissue barrier integrity of the respiratory epithelium. Overall, ACE2 expression was higher in the COPD than in the healthy epithelium (Figures 2 and S1-2), which is likely due to the presence of goblet cell hyperplasia in the COPD epithelium and thus a higher number of goblet cells.

TMPRSS2 is an important host co-factor for SARS-CoV-2 entry into target cells (Hoffmann et al., 2020; Lukassen et al., 2020; Shulla et al., 2011). We visualized TMPRSS2 in the apical site of the airway epithelium by staining with anti-TMPRSS2 and found that TMPRSS2 expression hardly overlapped with cilium (Figure 3A). We then tested the expression of TMPRSS2 along with that of the goblet cell markers MUC5AC and MUC5B. Indeed, TMPRSS2 overlapped with MUC5AC (Figure 3B) and MUC5B (Figures 3C). Therefore, it appears that both TMPRSS2 and ACE2 are expressed on the same cell surface (Figures 2 and 3). These results indicate that goblet cells may be a novel target of SARS-CoV-2 infection in the respiratory epithelium.

**Figure 3.**
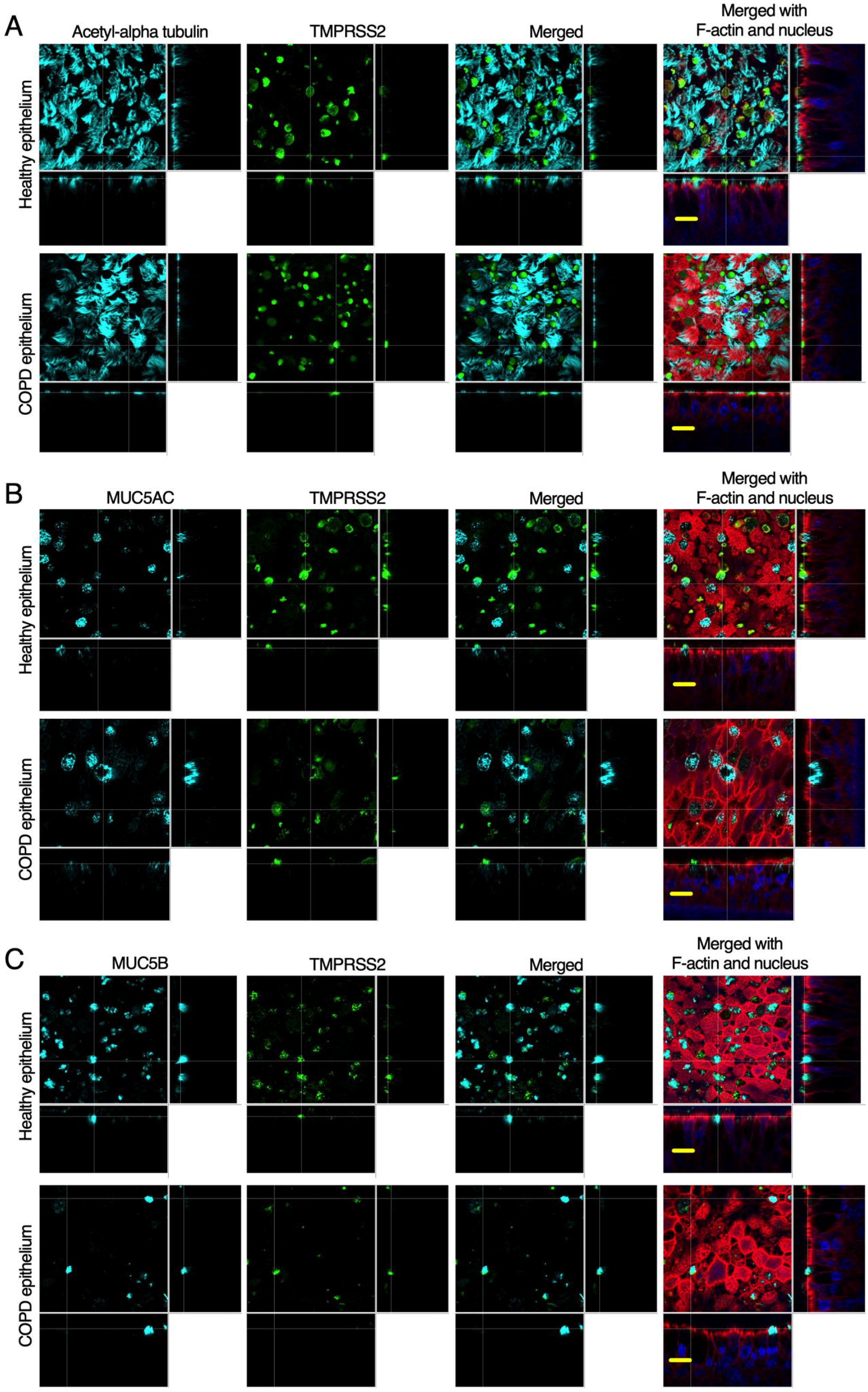
TMPRSS2 is expressed at higher levels in goblet cells. **(A).** The airway epithelia (described in Figure 1) were stained for cilia (anti-acetyl-alpha tubulin, cyan), TMPRSS2 (anti-TMPRSS2, green), and F-actin (rhodamine phalloidin, red); the nuclei were stained with DAPI (blue). **(B)**. The airway epithelia were stained for the identification of MUC5AC^+^ goblet cells (anti-MUC5AC, cyan), TMPRSS2 (anti-TMPRSS2, green), F-actin (rhodamine phalloidin, red) and nuclei (DAPI, blue). **(C).** The airway epithelia were stained to identify MUC5B^+^ goblet cells (anti-MUC5B, cyan), TMPRSS2 (anti-TMPRSS2, green), F-actin (rhodamine phalloidin, red) and nuclei (DAPI, blue). Bar =10 μm.

### SARS-CoV-2 infects goblet cells

Although we confirmed that SARS-CoV-2 entry receptors are expressed at higher levels on nonciliated goblet cells, a number of previous studies have suggested that SARS-CoV-2 targets ciliated cells (Hou et al., 2020; Lamers et al., 2020). Therefore, we first examined whether SARS-CoV-2 infects nonciliated goblet cells. We infected the apical side of the airway epithelium with SARS-CoV-2 at a multiplicity of infection of 0.1 (MOI = 0.1). At 4 days post infection (DPI), we fixed the cells and stained them for SARS-CoV-2 nucleoprotein (N) and a ciliated cell marker. The results revealed that SARS-CoV-2 infects both healthy and COPD epithelium and causes a substantial cytopathic effect (CPE) (Figures S3A and B). SARS-CoV-2 infection was higher in the COPD epithelium than in the healthy epithelium, as will be addressed later in the manuscript. Although in some cases the virus-induced extensive CPE made it difficult to distinguish SARS-CoV-2 cell tropism, we focused on multiple random areas with less CPE but virus infection. We found that SARS-CoV-2 infects nonciliated cells in both healthy and COPD epithelium (Figure 4A). We also used a second detection method to visualize viral and cellular markers in cross sections of the epithelium. Immunohistochemistry-based staining confirmed the extensive CPE induced by SARS-CoV-2 in both healthy and COPD epithelium. Apparently, SARS-CoV-2 infected both ciliated and nonciliated epithelial cells in the airway epithelium (Figure 4B). Using staining strategies similar to those described before, we found that SARS-CoV-2 infects MUC5B-positive (Figures 4C and D) and MUC5AC-postive (Figure 4E) goblet cells. To determine whether SARS-CoV-2 infects basal cells, we stained the sectioned epithelium for P63 and the SARS-CoV-2 spike (S) protein. We did not observe any overlap between the SARS-CoV-2 S protein and the basal cell marker (Figure S4). These results suggest that SARS-CoV-2 infects nonciliated goblet cells in addition to ciliated cells.

**Figure 4.**
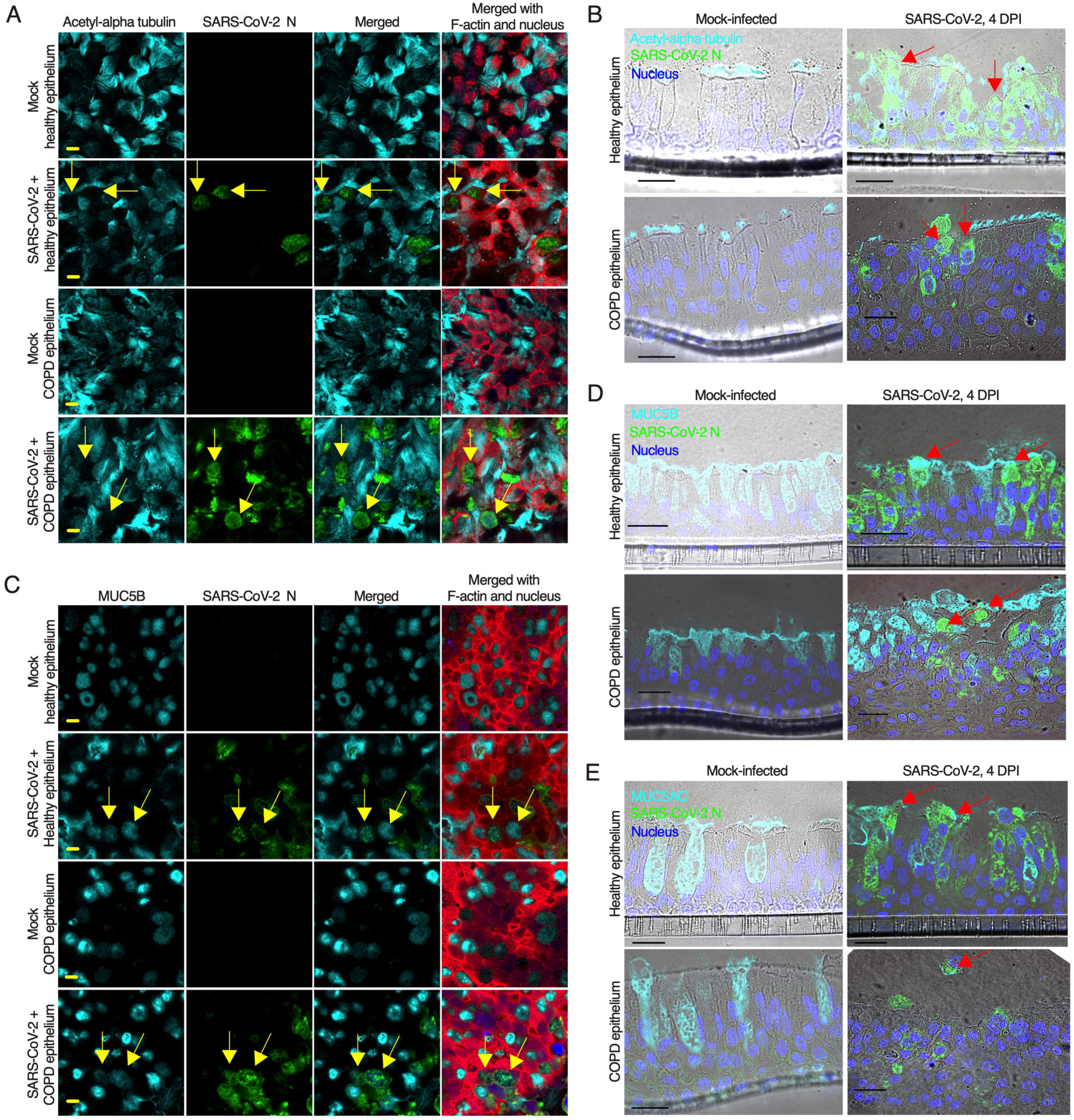
SARS-CoV-2 infects goblet cells. **(A).** The airway epithelia were mock-infected or infected with SARS-CoV-2 at an MOI of 0.1. At 4 days post infection (DPI), the epithelia were fixed, permeabilized, and stained for the identification of cilia (anti-acetyl-alpha tubulin, cyan), SARS-CoV-2 N (anti-N, green), F-actin (rhodamine phalloidin, red) and nuclei (DAPI, blue). The arrows indicate SARS-CoV-2-infected nonciliated cells. **(B).** At 4 DPI, the epithelia (described in A) were stained for the goblet cell marker MUC5B (anti-MUC5B, cyan), SARS-CoV-2 N (anti-N, green), and F-actin (rhodamine phalloidin, red); the nuclei were stained with DAPI (blue). The arrows indicate SARS-CoV-2-infected MUC5B^+^ goblet cells. **(C).** At 4 DPI, the epithelia (described in A) were embedded in paraffin, sectioned and stained for the detection of cilia (anti-acetyl-alpha tubulin, cyan) and SARS-CoV-2 N (anti-N, green); the nuclei were stained with DAPI (blue). The arrows indicate SARS-CoV-2-infected nonciliated cells. **(D).** At 4 DPI, the sectioned epithelia were stained for the goblet cell marker MUC5AC (anti-MUC5AC, cyan) and SARS-CoV-2 N (anti-N, green); the nuclei were also stained with DAPI (blue). The arrows indicate SARS-CoV-2-infected MUC5AC^+^ goblet cells. **(E).** At 4 DPI, the sectioned epithelia were stained for the goblet cell marker MUC5B (anti-MUC5B, cyan) and SARS-CoV-2 N (anti-N, green); the nuclei were also stained with DAPI (blue). The arrows indicate SARS-CoV-2-infected MUC5B^+^ goblet cells. (A and B) Bar = 5 μm. (C, D and E) Bar = 20 μm. See also Figures S3 and S4.

### SARS-CoV-2 induces syncytia and cell sloughing in the airway epithelium

To determine whether SARS-CoV-2 infection in the airway epithelium recapitulates the virus-induced pathophysiology in the lung, we examined the infected epithelium under a confocal microscope and found that SARS-CoV-2 infection causes substantial damage to the apical site of the infected epithelium, as confirmed by extensive CPE in both healthy and COPD epithelium (Figure S3). We also found substantial mucus secretion due to SARS-CoV-2 infection. The apical damage of the SARS-CoV-2-infected epithelium included loss of cellular junctions, loss of ciliary damage, substantial mucus production, and the protraction of nuclei, which are all common features observed in SARS-CoV-2-infected lungs (Schaefer et al., 2020). Additionally, we investigated whether SARS-CoV-2-infected cells in the epithelium might form syncytium (multinucleated cell), a hallmark of SARS-CoV-2 infection in the lung (Giacca et al., 2020) that had also been reported for SARS-CoV-1 (Franks et al., 2003). Indeed, we found that SARS-CoV-2-infected cells formed syncytia in both healthy and COPD epithelia (Figure 5A and B). Cell sloughing has been reported from lung autopsy findings of SARS-CoV-2-infected patients (Schaefer et al., 2020). Therefore, we examined whether SARS-CoV-2 infection in our epithelium model recapitulates cell sloughing. Indeed, we found that SARS-CoV-2 induces cell sloughing in both healthy and COPD epithelia as confirmed by two independent methods, immunofluorescence and IHC (Figures 5C and D). These results demonstrate that hallmark pathological features of SARS-CoV-2 are recapitulated in the infected airway epithelium model.

**Figure 5.**
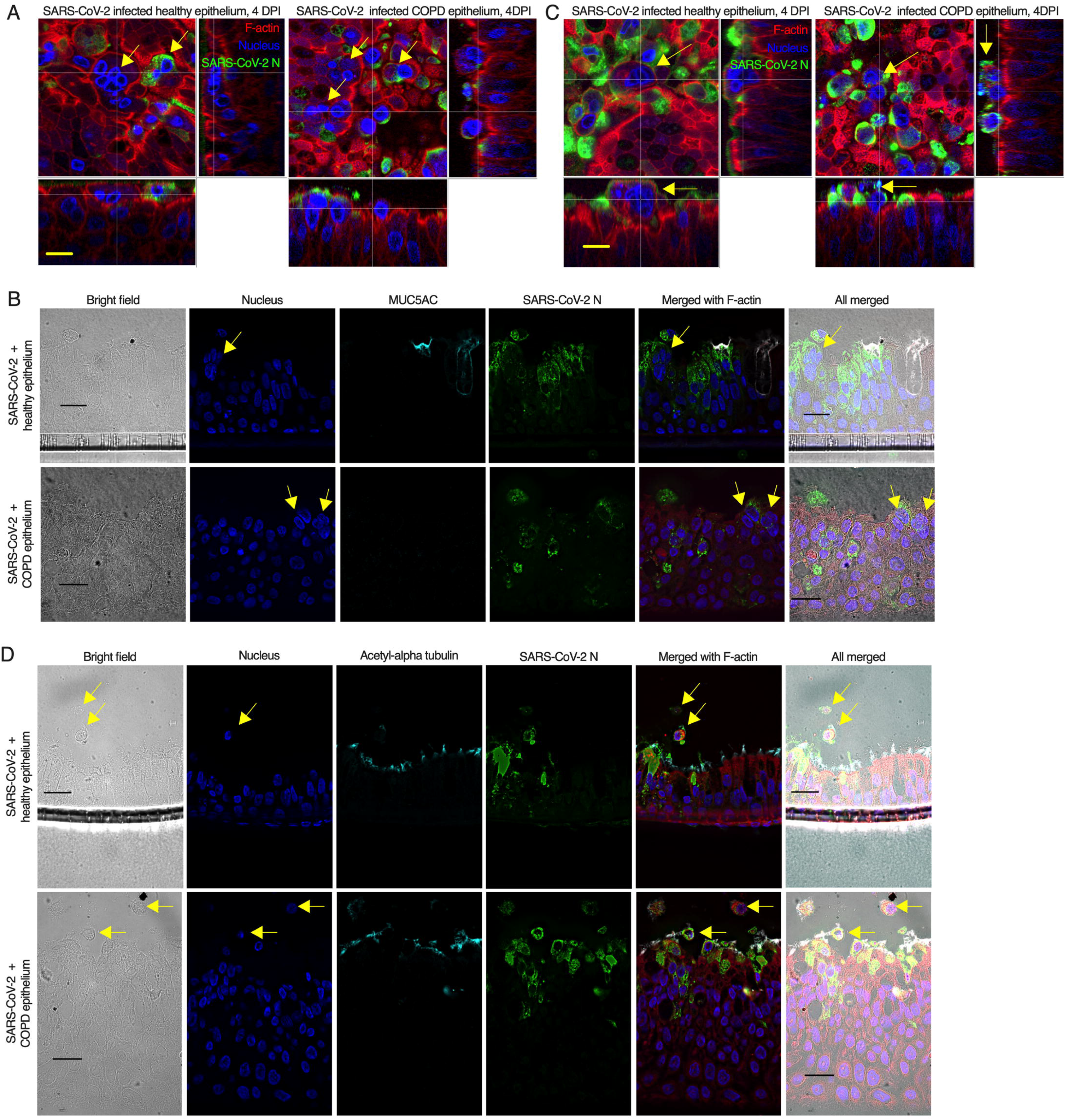
SARS-CoV-2 induces syncytia and cell sloughing. **(A).** The airway epithelia (mock or infected; described in Figure 4) were stained for SARS-CoV-2 N (anti-N, green) and F-actin (rhodamine phalloidin, red); the nuclei were also stained with DAPI (blue). The arrows indicate syncytia. Bar = 5 μm. **(B).** At 4 DPI, the sectioned epithelia were stained for the goblet cell marker MUC5AC (anti-MUC5AC, cyan) and SARS-CoV-2 N (anti-N, green), and the nuclei were also stained with DAPI (blue). The arrows indicate syncytia. Bar = 20 μm. **(C).** At 4 DPI, the airway epithelia were stained for SARS-CoV-2 N (anti-N, green) and F-actin (rhodamine phalloidin, red), and the nuclei were stained with DAPI (blue). The arrows indicate cell sloughing. Bar = 5 μm. **(D).** At 4 DPI, the sectioned epithelia were stained for cilia (anti-acetyl-alpha tubulin, cyan) and SARS-CoV-2 N (anti-N, green), and the nuclei were stained with DAPI (blue). The arrows indicate cell sloughing. Bar = 20 μm.

### SARS-CoV-2 replicates higher and exacerbates pathophysiology in COPD epithelium

To determine whether SARS-CoV-2 replicates better in the COPD epithelium, we titered SARS-CoV-2 in the apical wash of the infected epithelium and found that SARS-CoV-2 replication was increased by almost a log in the COPD compared to the healthy epithelium (Figure 6A). In addition, we found a squamous metaplasia in SARS-CoV-2-infected COPD epithelium which was rather infrequently found in the SARS-CoV-2-infected healthy epithelium (Figures 1I; 4B, D, and E, and 5B and D) (Borczuk et al., 2020). Squamous metaplasia is known to increase bronchial wall thickening as seen in bronchitis (Randell, 2006; Reid et al., 2018; Rigden et al., 2016). As tracheobronchitis is one of the most common histopathological features in the COVID-19 disease fatalities (Martines et al., 2020), we evaluated whether SARS-CoV-2 infection increases height of the epithelium. First, there was a substantial increase in squamous metaplasia in the COPD epithelium due to SARS-CoV-2 infection (Figure S5). Second, the increased metaplasia apparently changed the morphology of the nonciliated goblet cells in the infected COPD epithelium (Figures 6B and C). Whether this change in the goblet cell morphology impacted the mucus hyperplasia remains to be determined. Third, in contrast to the healthy epithelium, SARS-CoV-2 induced higher squamous metaplasia in the infected COPD epithelium and caused a substantial increase in the height of the epithelium (Figure 6D). These results suggest that SARS-CoV-2 replicates better in the COPD epithelium and exacerbates pathophysiology in the infected airway epithelium.

**Figure 6.**
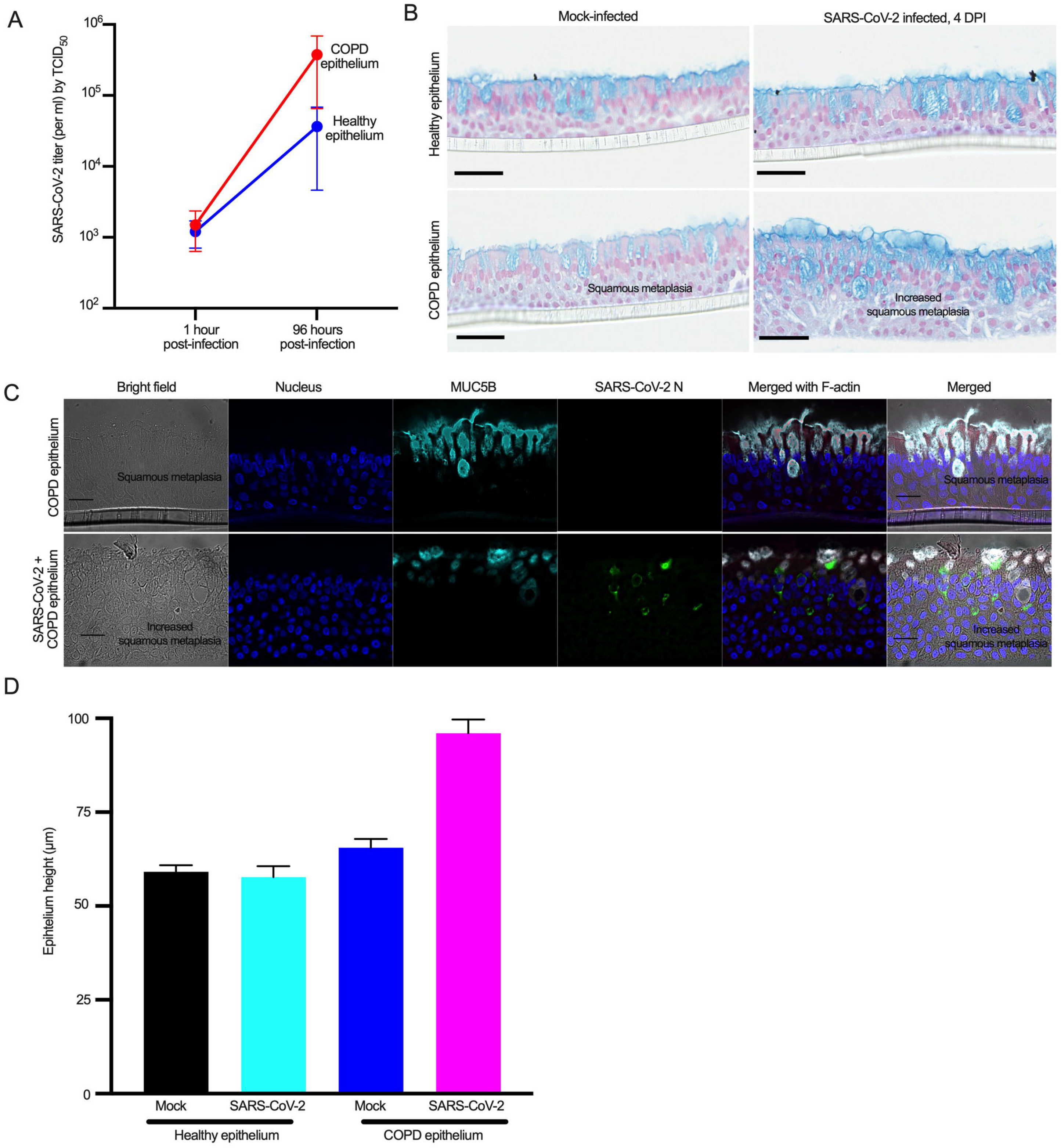
SARS-CoV-2 replicates at a higher rate and increases squamous metaplasia in the COPD epithelium. **(A).** The apical wash of the SARS-CoV-2-infected airway epithelia (described in Figure 4) was collected, and SARS-CoV-2 titration was performed based on the tissue culture infective dose 50 (TCID_50_). The results from an independent experiment (n=3) are shown. The error bars show the SEMs. **(B).** The sectioned epithelia (described in Figure 4) were stained with Alcian blue. Bar = 50 μm. **(C).** At 4 DPI, the sectioned epithelia (COPD epithelia after mock or SARS-CoV-2 infection) were stained for the goblet cell marker MUC5B (anti-MUC5B, cyan) and SARS-CoV-2 N (anti-N, green), and the nuclei were stained with DAPI (blue). Bar = 20 μm. **(D).** The height of the airway epithelia (described in C) was measured from the images (n=5), the average of 12 points was plotted. See also Figure S5.

## Discussion

Here, we have shown that the COPD epithelium model recapitulates the bronchial biophysical and pathophysiological characteristics of COPD, such as goblet cell hyperplasia and squamous metaplasia (Gohy et al., 2019; Kim et al., 2015; Reid et al., 2018; Rigden et al., 2016). A previous report suggested the presence of altered ciliated cells in COPD airway epithelium (Gohy et al., 2019), which was not observed in our model. The duration of cell differentiation may be the reason, as we differentiated NHBE cells for four weeks that may require to see terminal differentiation (including ciliogenesis) (Gohy et al., 2019). We used an air-liquid interface (ALI) culture method to generate multi-cellular diversity and physiologic functioning airway epithelium that resembles the airway surface in vivo (Fulcher et al., 2005; Pawlina, 2016; Rayner et al., 2019). One of the limitations in the ALI culture research is passaging of primary NHBE cells may impact on their ability to differentiate into airway epithelium. We could demonstrate that primary NHBE cells obtained after four passages without using any additional supplements can still be differentiated into human airway epithelium (Rayner et al., 2019). In a separate study, we confirmed that passaging NHBE cells up to four times has insignificant effect on the whole-genome transcriptome by comparing transcriptome profiles of each passage cells (data not shown). The ability to expand primary cells that also form fully differentiated mucociliary epithelium reduces repeat sample collections from patients where samples are difficult and limited, such as infants and deceased patients (Rayner et al., 2019; Wolf et al., 2017). Our results demonstrate primary NHBE cells either from healthy or a COPD patient can be passaged up to four times and that normal epithelial phenotypic features are maintained in passaged primary NHBE cells.

One emergent question is why the human-to-human transmission of SARS-CoV-2 is much higher compared to SARS-CoV-1, although both viruses share ACE2 as cell surface receptor and use TMPRSS2 to facilitate their entry into the host cell (Hoffmann et al., 2020; Lukassen et al., 2020). The SARS-CoV-2 spike protein has an additional furin cleavage site that is absent in SARS-CoV-1, and it is hypothesized that furin cleavage facilitates human-to-human transmission (Coutard et al., 2020; Lukassen et al., 2020). We found that SARS-CoV-2 infected both ciliated cells and nonciliated goblet cells, but not basal cells in the airway epithelium. Although SARS-CoV-2 may preferentially infect goblet cells due to the higher expression of ACE2 and TMPRSS2, further studies are needed to confirm temporal and spatial regulations of SARS-CoV-2 infection in the airway epithelium. The major function of goblet cells in the lung epithelium is mucin production to trap pathogens, dust, and particles, which are cleared by a process known as mucociliary clearance (Rogers, 2003). The possibility that SARS-CoV-2 infects goblet cells could explain the presence of viral RNA in sputum (Wang et al., 2020) and might explain the easy transmission of the virus from person to person. While we are preparing our manuscript, Hao et al., have shown that goblet cells are permissive to SARS-CoV-2 infection (Hao et al., 2020). As SARS-CoV-1 infection is limited to ciliated cells (Sims et al., 2005), we think that SARS-CoV-2 infection in goblet cells could explain why SARS-CoV-2 is more transmissible than SARS-CoV-1. Influenza A virus infects goblet cells in addition to ciliated cells (Matrosovich et al., 2004), whereas respiratory syncytial virus (RSV) infects only ciliated cells (Zhang et al., 2002). Therefore, virus-specific preferential cell tropism in the lung could explain the difference in respiratory virus pathogenicity and transmissibility.

Our airway epithelium model recapitulated some SARS-CoV-2-induced lung histopathological findings, such as prominent nucleoli (Schaefer et al., 2020), cytoplasmic vacuolation, multinucleated giant cells (Falasca et al., 2020), squamous metaplasia (Martines et al., 2020) and epithelial cell sloughing, which is the most common finding (Bradley et al., 2020). SARS-CoV-2 infection in the bronchial epithelium models induced a substantial amount of cell syncytium formation and cell sloughing, which indicates that tracheobronchial cells are highly associated with virus-induced pathogenesis. Therefore, our data suggest that bronchial cells are equally important to alveolar cells in the SARS-CoV-2 pathogenesis (Martines et al., 2020). In patients with COPD, goblet cell hyperplasia is a common clinical manifestation, and an increasing number of goblet cells results in more mucus production (Shaykhiev, 2019). Because we have shown that SARS-CoV-2 replicated at a higher rate in the COPD epithelium, we believe that goblet cell hyperplasia is responsible for this phenomenon and may explain why COPD patients are at higher risk for severe outcome of COVID-19. Interestingly, we found that SARS-CoV-2 infection induced “squamous metaplasia” in the COPD epithelium. Squamous metaplasia in the columnar epithelium corelates with increased severity of airway obstruction in COPD (Araya et al., 2007). Although squamous metaplasia is one of the histopathological findings in the SARS-CoV-2 infected patient fatalities (Martines et al., 2020), our results provide evidence for the first time that SARS-CoV-2 infection increases squamous metaplasia, which may result in bronchitis in the infected COPD patients. In fact, tracheobronchitis is one of the most common histopathological features in the COVID-19 disease fatalities (Martines et al., 2020). No SARS-CoV-2 infectious virion was detected in the basal medium of the infected healthy and COPD epithelia, which might suggest that the initial virus-induced substantial damage was limited at the apical site.

Immune responses to viral infection play a critical role in determining the clinical outcomes of SARS-CoV-2 infection (Huang et al., 2020). Unfortunately, our airway epithelium model lacks both resident and infiltrating immune cells (dendritic cells, macrophages, and lymphocytes, etc.) and is devoid of the endothelial layer, and therefore cannot study the impact of immune responses on the SARS-CoV-2 infections. However, we can provide inferences based on the detection of immunoregulatory factors (chemokines and cytokines), but these hypotheses remain to be confirmed. Overall, we believe that the airway epithelium model provides an excellent tool for demystifying some of the SARS-CoV-2 pathophysiological features and identifying and testing novel therapeutics against the virus.

In conclusion, we developed in vitro lung airway epithelium models from passaged primary NHBE cells of either a healthy or a COPD patient. Our studies revealed SARS-CoV-2 infection of goblet cells leading to virus-induced syncytium formation and cell sloughing in the airway epithelium. We found that SARS-CoV-2 replicates better COPD airway epithelium likely due to COPD associated goblet cell hyperplasia. Thus, we postulate that goblet cells play a critical role in SARS-CoV-2 infection of the lung and are responsible for more severe outcome of SARS-CoV-2 infection in COPD patients.

## Graphical Abstract

### Highlights

1. The airway epithelium model recapitulates many bronchial pathological characteristics of COPD, such as goblet cell hyperplasia and squamous metaplasia
2. ACE2 and TMPRSS2 are highly expressed on nonciliated goblet cells
3. SARS-CoV-2 infects goblet cells and induces syncytium formation and cell sloughing
4. SARS-CoV-2 replicates at a higher rate in the COPD epithelium and increases squamous metaplasia

### In Brief

Osan et al. showed that SARS-CoV-2 preferentially infects and replicates in nonciliated goblet cells inducing syncytium formation and cell sloughing. Our results suggest that goblet cells play a critical role in SARS-CoV-2-induced pathophysiology in the lung.

## Acknowledgments

We thank the Microscopy Core (UND, Grand Forks) funded by NIH P20GM103442 of the INBRE program for providing access to an Olympus FV300 confocal microscope and Dr. Swojani Shrestha for the technical support provided. Histological services were provided by Histology Core (UND, Grand Forks) funded by NIH P20GM113123, NIH DaCCoTA CTR, NIH U54GM128729, and UNDSMHS funds. We also thank the Imaging Core (UND, Grand Forks) funded by NIH P20GM113123, NIH U54GM128729, and UNDSMHS funds for the IMARIS image analyses. We thank John Lee, Information Resources, UND, for preparing the graphical abstract. This work was funded by NIH/NIGMS P20GM113123 and in part by the Intramural Research Program of NIAID, NIH.

## Author contributions

M.M., J.K.O., and S.N.T. conceived the project and designed the experiments. K.L.B. provided the primary cells as well as training and guidance on primary cell differentiation. M.M., J.K.O., and S.N.T. performed the experiments. S.N.T. generated the confocal images. F.F. and H.F. performed the virus infection experiments in the BSL-4 laboratory. J.K.O., B.A.D., and K.J. generated the IHC images, B.A.D. performed the H&E and Alcian Blue staining, and M.M., J.K.O., and S.N.T. wrote the paper. M.M. and H.F. reviewed and edited the paper.

## STAR METHOD

### KEY RESOURCE TABLE

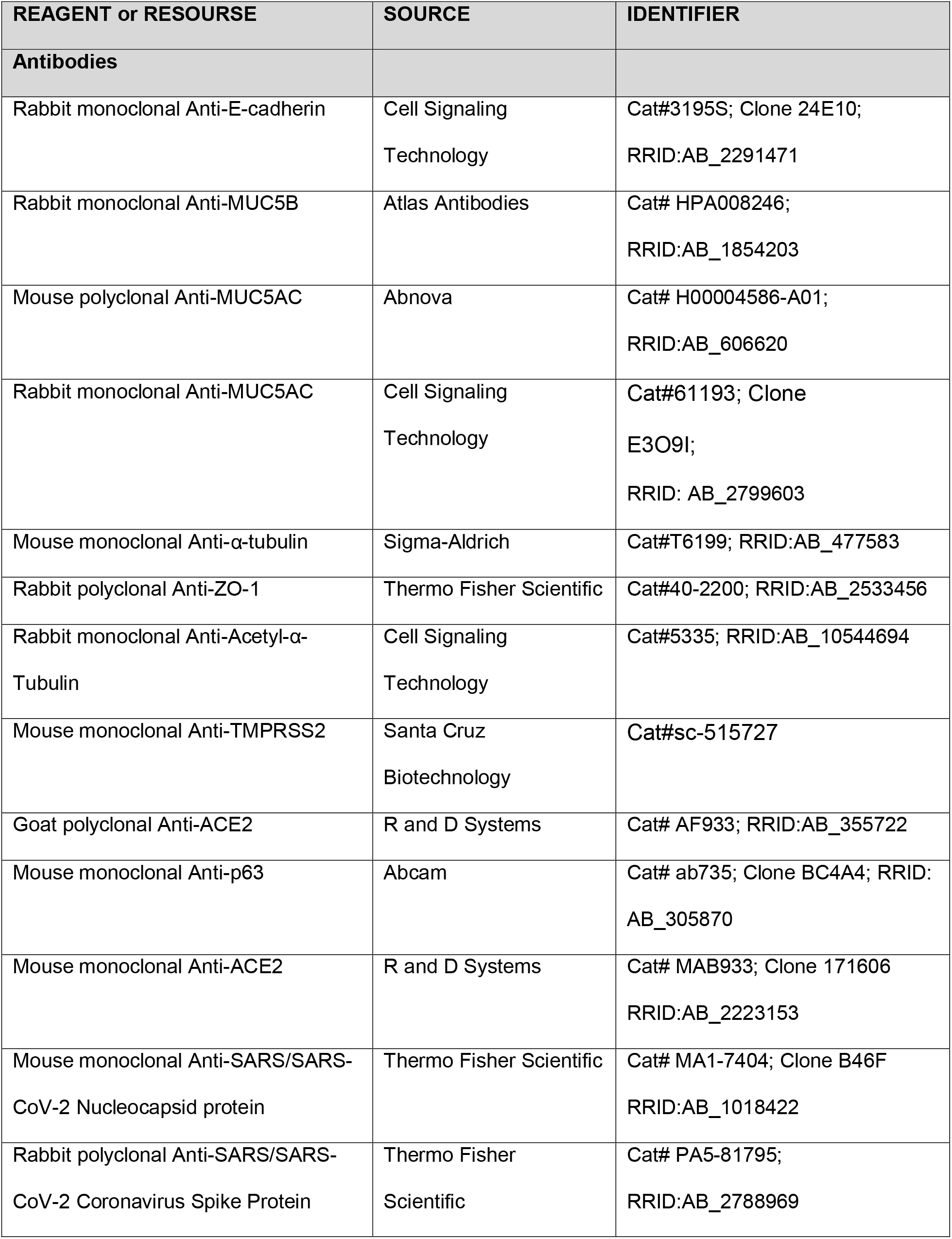
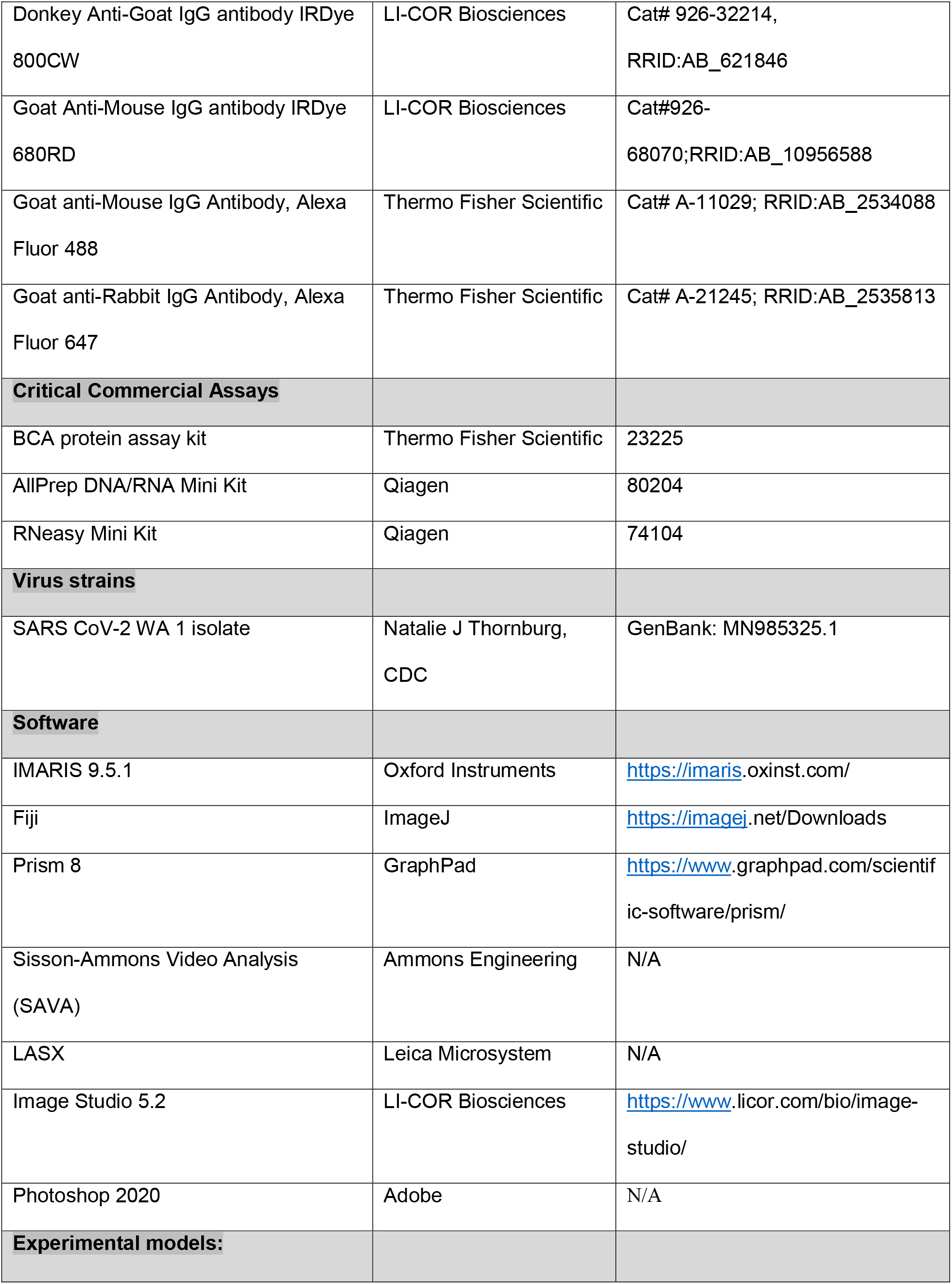
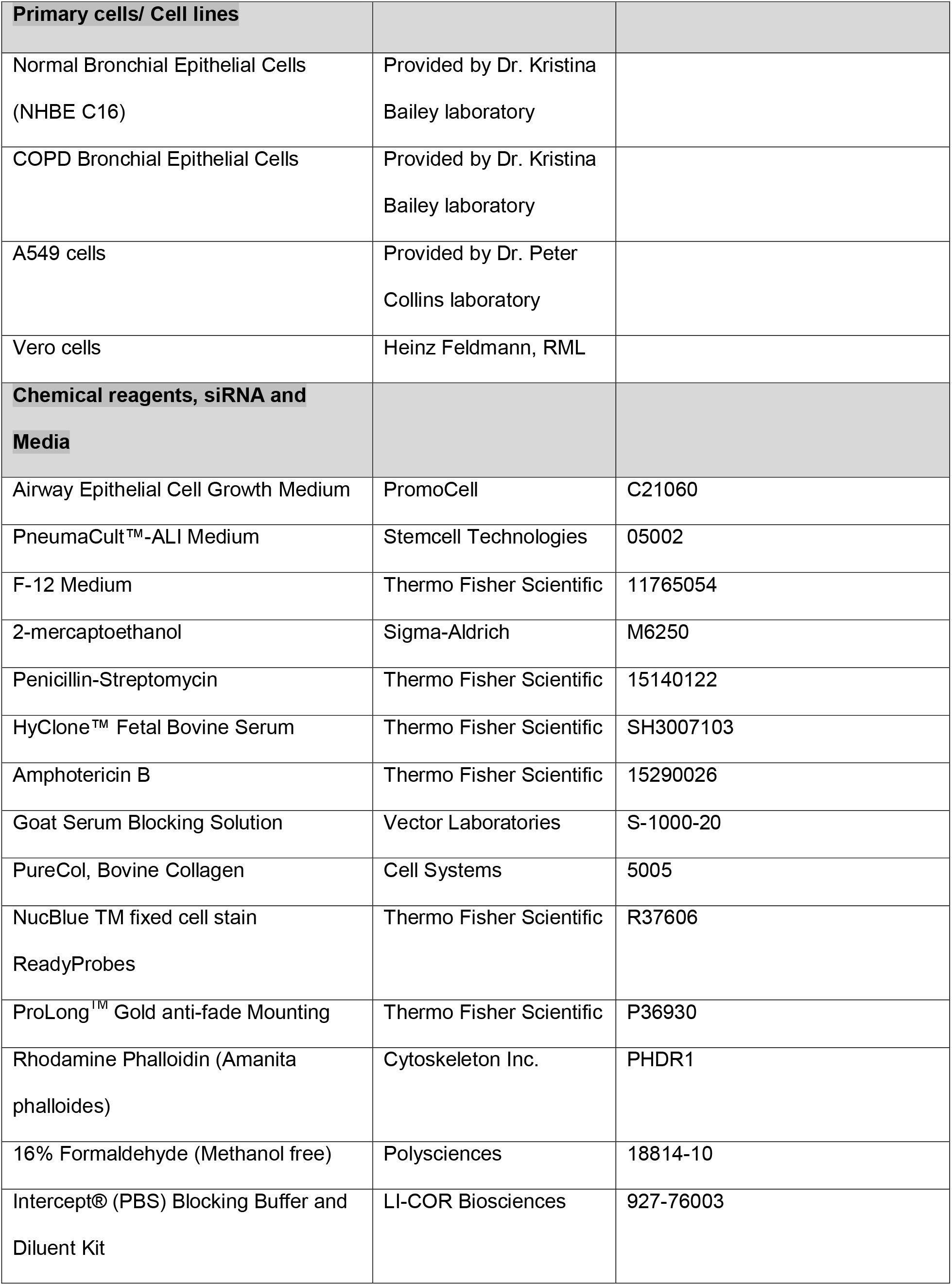
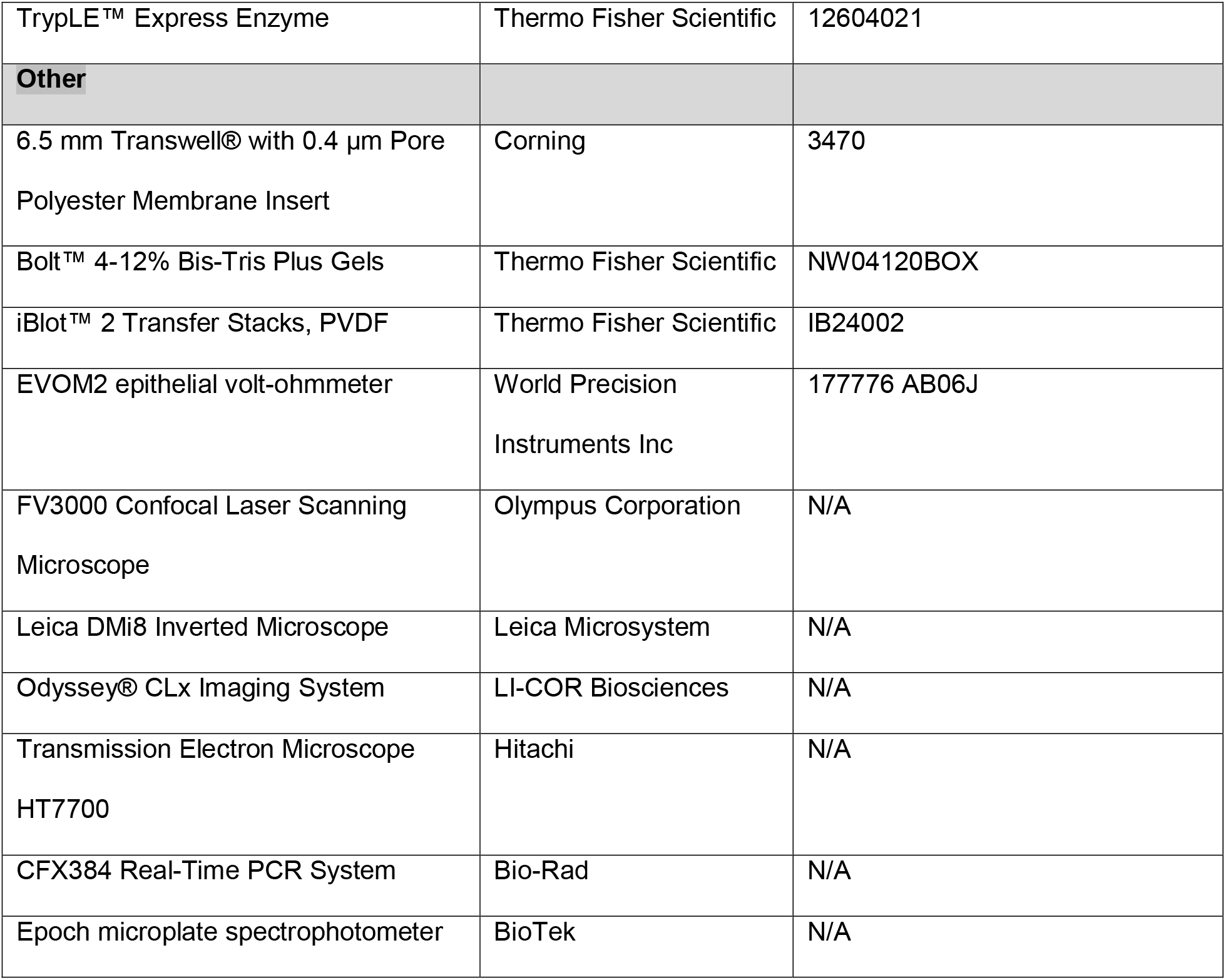

**STAR METHOD**
| REAGENT or RESOURCE | SOURCE | IDENTIFIER |
| --- | --- | --- |
| **Antibodies** |  |  |
| Rabbit monoclonal Anti-E-cadherin | Cell Signaling Technology | Cat#3195S; Clone 24E10;<br>RRID:AB_2291471 |
| Rabbit monoclonal Anti-MUC5B | Atlas Antibodies | Cat# HPA008246;<br>RRID:AB_1854203 |
| Mouse polyclonal Anti-MUC5AC | Abnova | Cat# H00004586-A01;<br>RRID:AB_606620 |
| Rabbit monoclonal Anti-MUC5AC | Cell Signaling Technology | Cat#61193; Clone E3O9I;<br>RRID: AB_2799603 |
| Mouse monoclonal Anti-α-tubulin | Sigma-Aldrich | Cat#T6199; RRID:AB_477583 |
| Rabbit polyclonal Anti-ZO-1 | Thermo Fisher Scientific | Cat#40-2200; RRID:AB_2533456 |
| Rabbit monoclonal Anti-Acetyl-α-Tubulin | Cell Signaling Technology | Cat#5335; RRID:AB_10544694 |
| Mouse monoclonal Anti-TMPRSS2 | Santa Cruz Biotechnology | Cat#sc-515727 |
| Goat polyclonal Anti-ACE2 | R and D Systems | Cat# AF933; RRID:AB_355722 |
| Mouse monoclonal Anti-p63 | Abcam | Cat# ab735; Clone BC4A4; RRID: AB_305870 |
| Mouse monoclonal Anti-ACE2 | R and D Systems | Cat# MAB933; Clone 171606<br>RRID:AB_2223153 |
| Mouse monoclonal Anti-SARS/SARS-CoV-2 Nucleocapsid protein | Thermo Fisher Scientific | Cat# MA1-7404; Clone B46F<br>RRID:AB_1018422 |
| Rabbit polyclonal Anti-SARS/SARS-CoV-2 Coronavirus Spike Protein | Thermo Fisher Scientific | Cat# PA5-81795;<br>RRID:AB_2788969 |
| Donkey Anti-Goat IgG antibody IRDye 800CW | LI-COR Biosciences | Cat# 926-32214, RRID:AB_621846 |
| Goat Anti-Mouse IgG antibody IRDye 680RD | LI-COR Biosciences | Cat#926-68070;RRID:AB_10956588 |
| Goat anti-Mouse IgG Antibody, Alexa Fluor 488 | Thermo Fisher Scientific | Cat# A-11029; RRID:AB_2534088 |
| Goat anti-Rabbit IgG Antibody, Alexa Fluor 647 | Thermo Fisher Scientific | Cat# A-21245; RRID:AB_2535813 |
| **Critical Commercial Assays** |  |  |
| BCA protein assay kit | Thermo Fisher Scientific | 23225 |
| AllPrep DNA/RNA Mini Kit | Qiagen | 80204 |
| RNeasy Mini Kit | Qiagen | 74104 |
| **Virus strains** |  |  |
| SARS CoV-2 WA 1 isolate | Natalie J Thornburg, CDC | GenBank: MN985325.1 |
| **Software** |  |  |
| IMARIS 9.5.1 | Oxford Instruments | https://imaris.oxinst.com/ |
| Fiji | ImageJ | https://imagej.net/Downloads |
| Prism 8 | GraphPad | https://www.graphpad.com/scientific-software/prism/ |
| Sisson-Ammons Video Analysis (SAVA) | Ammons Engineering | N/A |
| LASX | Leica Microsystem | N/A |
| Image Studio 5.2 | LI-COR Biosciences | https://www.licor.com/bio/image-studio/ |
| Photoshop 2020 | Adobe | N/A |
| **Experimental models:** |  |  |

| **Primary cells/ Cell lines** |  |  |
| --- | --- | --- |
| Normal Bronchial Epithelial Cells<br>(NHBE C16) | Provided by Dr. Kristina<br>Bailey laboratory |  |
| COPD Bronchial Epithelial Cells | Provided by Dr. Kristina<br>Bailey laboratory |  |
| A549 cells | Provided by Dr. Peter<br>Collins laboratory |  |
| Vero cells | Heinz Feldmann, RML |  |
| **Chemical reagents, siRNA and**<br>**Media** |  |  |
| Airway Epithelial Cell Growth Medium | PromoCell | C21060 |
| PneumaCult™-ALI Medium | Stemcell Technologies | 05002 |
| F-12 Medium | Thermo Fisher Scientific | 11765054 |
| 2-mercaptoethanol | Sigma-Aldrich | M6250 |
| Penicillin-Streptomycin | Thermo Fisher Scientific | 15140122 |
| HyClone™ Fetal Bovine Serum | Thermo Fisher Scientific | SH3007103 |
| Amphotericin B | Thermo Fisher Scientific | 15290026 |
| Goat Serum Blocking Solution | Vector Laboratories | S-1000-20 |
| PureCol, Bovine Collagen | Cell Systems | 5005 |
| NucBlue TM fixed cell stain<br>ReadyProbes | Thermo Fisher Scientific | R37606 |
| ProLong™ Gold anti-fade Mounting | Thermo Fisher Scientific | P36930 |
| Rhodamine Phalloidin (Amanita<br>phalloides) | Cytoskeleton Inc. | PHDR1 |
| 16% Formaldehyde (Methanol free) | Polysciences | 18814-10 |
| Intercept® (PBS) Blocking Buffer and<br>Diluent Kit | LI-COR Biosciences | 927-76003 |
| TrypLE™ Express Enzyme | Thermo Fisher Scientific | 12604021 |
| **Other** |  |  |
| 6.5 mm Transwell® with 0.4 µm Pore Polyester Membrane Insert | Corning | 3470 |
| Bolt™ 4-12% Bis-Tris Plus Gels | Thermo Fisher Scientific | NW04120BOX |
| iBlot™ 2 Transfer Stacks, PVDF | Thermo Fisher Scientific | IB24002 |
| EVOM2 epithelial volt-ohmmeter | World Precision Instruments Inc | 177776 AB06J |
| FV3000 Confocal Laser Scanning Microscope | Olympus Corporation | N/A |
| Leica DMI8 Inverted Microscope | Leica Microsystem | N/A |
| Odyssey® CLx Imaging System | LI-COR Biosciences | N/A |
| Transmission Electron Microscope HT7700 | Hitachi | N/A |
| CFX384 Real-Time PCR System | Bio-Rad | N/A |
| Epoch microplate spectrophotometer | BioTek | N/A |

## RESOURCE AVIALABILITY

### Lead Contact

Further information and requests for resources and reagents should be directed to and will be fulfilled by the lead contact, Masfique Mehedi.

### Material Availability

The materials and reagents generated in this study will be made available upon installment of a material transfer agreement (MTA).

## EXPERIMENTAL MODEL AND SUBJECT DETAILS

### Cells

Primary normal human bronchial epithelial (NHBE) cells from either a deidentified healthy adult (female, aged 52 years, never-smoker) or a patient with chronic obstructive pulmonary disease (COPD) were obtained under an approved material transfer agreement (MTA) between the Mehedi Laboratory at University of North Dakota (UND) and Bailey Laboratory at University of Nebraska Medical Center (UNMC), Omaha, NE. A549 cells (a human lung epithelial cell line, ATCC-185) were obtained from Dr. Peter Collins at the National Institutes of Health (NIH), Bethesda, MD, USA. Vero cells used for TCID_50_ assay were a resource of the Feldmann Lab at the Rocky Mountain Laboratories, Hamilton, MT.

### Virus

The SARS-CoV-2 isolate nCoV-WA1-2020 (MN985325.1) was kindly provided by CDC as Vero passage 3 (Harcourt et al., 2020). The virus was propagated once in Vero E6 cells in DMEM (Sigma) supplemented with 2% fetal bovine serum (Thermo Fisher Scientific), 1 mM L-glutamine (Thermo Fisher Scientific), 50 U/ml penicillin and 50 μg/ml streptomycin (Thermo Fisher Scientific) (virus isolation medium). The virus stock used in this study was 100% identical to the initially deposited GenBank sequence (MN985325.1); sequencing did not detect any virus stock contaminants.

### Biosafety Statement

Work with SARS-CoV-2 was performed in the high biocontainment facilities at the Rocky Mountain Laboratories (RML), NAID, NIH in Hamilton, MT. All infectious work followed standard operating procedures (SOPs) approved by the Institutional Biosafety Committee.

## METHOD DETAILS

### Cell culture

We passaged the primary NHBE cells (passage 1) three times before differentiating them (passage 4) into a pseudostratified epithelium. For each passage, the cells were grown in a 100-mm culture dish (Corning Inc.) precoated with PurCol (Advanced Biometrics). The cells were maintained in airway epithelial cell (AEC) growth medium (PromoCell) containing AEC supplements (PromoCell), 2% penicillin/streptomycin (Thermos Fisher Scientific), and 1% amphotericin B (Thermos Fisher Scientific) (complete AEC medium) at 37°C in an incubator with 5% CO_2_. The cells were grown to 90% confluency, and the medium was changed every other day. Confluent monolayers of cells were dissociated with 5 ml of TrypLE (Thermo Fisher Scientific) and pelleted, and one-third of the cells were reseeded in a culture dish containing complete AEC medium for passaging. A549 cells were grown in F-12 medium (Thermo Fisher Scientific) with 10% HyClone fetal bovine serum (GE Healthcare), 2% penicillin/streptomycin, and 1% amphotericin B.

### Air-liquid interface (ALI) culture

Transwells (6.5 mm) with 0.4-μm-pore polyester membrane inserts (Corning Inc.) were coated with PureCol for 20 minutes before cell seeding. NHBE cells (5×10^4) suspended in 200 μl of complete AEC medium were seeded in the apical part of a Transwell. Subsequently, 500 μl of complete AEC medium was added to the basal part of the Transwell. When the cells formed a confluent layer on the Transwell insert, the AEC medium was removed from the apical part, and PneumaCult-ALI basal medium (Stemcell Technologies) with the required supplements (Stemcell Technologies), 2% penicillin/streptomycin and 1% amphotericin B (complete ALI basal medium) was added to the basal part. The ALI medium in the basal part was changed every other day, and the apical surface was washed with 1x Dulbecco’s phosphate-buffered saline (DPBS) (Thermo Fisher Scientific) once per week initially but more frequently when more mucus was observed on later days. However, the thickness of the mucus was not determined. All the cells were differentiated for at least 4 weeks at 37°C in an incubator with 5% CO_2_.

### Transportation of Transwells containing airway epithelia

The differentiated airway epithelia for 22 days were transported in transportation medium at 4°C via overnight FedEx biological material shipment to the Rocky Mountain Laboratories (RML, Hamilton, MT, USA). The semisolid transportation medium was prepared by adding agarose to the complete AEC medium. Upon receipt, the Transwells containing airway epithelia were immediately transferred into new plates with complete AEC medium in the basal part and maintained before infection as described in the section on ALI culture.

### Virus infection

Virus infection was conducted in the high biocontainment facilities at RML, NIAID, NIH. The 4-week-differentiated airway epithelia were washed with 200 μl of 1x PBS to remove mucus and infected on the apical site with SARS-CoV-2 at an MOI of 0.1 in 1x PBS for 1 hour (at 37°C with 5% CO_2_). For mock infection, the assigned Transwells were similarly incubated with 1x PBS without virus. The viral inoculum was then removed, and the epithelia were washed twice with 200 μl of 1x PBS. Two hundred microliters of apical wash and basal medium were collected for virus titration. Fresh ALI medium (1000 μl) with supplements was added to the basal part of each Transwell, and the apical part was kept empty. Mock-infected and SARS-CoV-2-infected Transwells were incubated for 4 days at 37°C in an incubator with 5% CO_2_.

### Tissue culture infective dose 50 (TCID_50_)

Vero cells were seeded on 96-well plates to confluency, and SARS-CoV-2 endpoint titration was performed. The cells were inoculated with 10-fold dilutions of the supernatants collected during the experiment in DMEM, 2% FBS, L-glutamine and P/S. The cells were incubated for 7 days, and the cytopathic effect (CPE) was scored under a microscope. The TCID_50_ was calculated using the Reed-Muench formula (Reed and Muench, 1938).

### Virus-infected sample collection

At 4 DPI, 200 μl of 1x PBS was added to the apical site of the Transwell and incubated for several minutes, and the apical wash was collected for virus titration. Similarly, basal medium was collected for virus titration. All the samples were stored at −80°C until titration. For PFA fixation, 200 μl of 4% PFA was added at the apical site of the Transwells and incubated for 30 minutes, and the Transwells were further maintained overnight in 4% PFA before being transported outside of the BSL4 lab according to the standard protocol of RML. For protein and RNA sample collections, 200 μl of 1x PBS was added to the apical side of each Transwell, and cells were scrapped with a mini scrapper, collected into a cryovial, and pelleted by being spun down with a table-top bench centrifuge. For the protein samples, 100 μl of 2x SDS buffer was added to the cryovials containing the cell pellet, and the tube was boiled for 10 minutes. After a quick spin, the samples were transferred into new tubes and removed from the BSL4 laboratory using a standard RML protocol. For the RNA samples, the cell pellets were resuspended in 600 μl of RLT buffer (Qiagen) and transported out of the laboratory based on the RML protocol with 600 μl of ethanol. RNA was extracted using a Qiagen RNeasy mini kit (QIAGEN) and eluted in 50 μl of RNase-free water. All the samples were shipped to Mehedi Lab at UND using an appropriate overnight biological shipment procedure.

### Hematoxylin and eosin (H&E) staining

Four days post infection, NHBE-ALI and COPD-ALI mock and infected Transwells were fixed with 4% PFA for 24 hours. The membrane was cut off using a scalpel and embedded in paraffin mold. Five-micrometer sections were cut using a microtome. H&E staining was performed by two 5-minute Histo-Clear incubations followed by three 5-minute incubations with 100% ethanol, one 5-minute incubation with 95% ethanol and one 5-minute incubation with 70% ethanol. Subsequently, the samples were washed with tap water for 2 minutes, incubated with 4% acidified Harries Hematoxylin for 2 minutes, washed with tap water for 2 minutes, and incubated with 0.5% lithium carbonate for 20 seconds. The samples were subsequently washed with tap water for 5 minutes, incubated with Eosin Y Solution with Pholoxine for 2 minutes and then subjected to a series of ethanol incubations: incubation with 70% ethanol for 5 minutes, incubation with 95% ethanol for 5 minutes and three 5-minutes incubation with 100% ethanol. Finally, two 5-minute incubations with Histo-Clear were performed, and coverslips were then placed using permanent mounting medium.

### Alcian blue staining

The 5-μm sectioned ALI-NHBE and COPD mock and infected samples were also subjected to Alcian blue staining. The sections were deparaffinized as described in the section on H&E staining using Histo-Clear and ethanol. The cells were then washed with tap water for 2 minutes, incubated in 3% acetic acid for 3 minutes and stained with Alcian blue for 30 minutes. The sections were subsequently rinsed with 3% acetic acid for 10 seconds to help prevent nonspecific staining. The sections were washed under tap water for 10 minutes and then rinsed for 2 minutes in Milli-Q water. The sections were counterstained with nuclear fast red solution for 5 minutes and washed under tap water for 5 minutes. The sections were subsequently dehydrated and cleared with a gradient of alcohol and Histo-Clear as in the protocol used for H&E staining. The slides were mounted with permanent mounting medium and coverslipped.

### Fluorescence imaging

The apical site of the airway epithelium was washed with PBS, and both the apical and basal parts were fixed with 4% paraformaldehyde (PFA) (Polysciences, Inc.), and blocked with 10% goat serum (Vector Laboratories) solution in immunofluorescence (IF) washing buffer (130 mM NaCl_2_, 7 mM Na_2_HPO_4_, 3.5 mM NaH_2_PO_4_, 7.7 mM NaN_3_, 0.1% BSA, 0.2% Triton-X 100 and 0.05% Tween-20) for 1 hour. The Transwell inserts were then incubated with the following primary antibodies (Abs) (alone or in combination) in IF washing buffer overnight at 4°C: anti-ZO-1 rabbit polyclonal (1:200) (Thermo Fisher Scientific), anti-E-cadherin rabbit monoclonal (1:200) (Cell Signaling Technologies), anti-MUC5B rabbit monoclonal (1:500) (Atlas Antibodies), anti-MUC5AC mouse polyclonal (1:100) (Abnova), anti-MUC5AC rabbit monoclonal (1:200) (Cell Signaling Technology), anti-acetyl-alpha-tubulin rabbit monoclonal (1:500) (Cell Signaling Technologies), anti-TMPRSS2 mouse monoclonal (1:50) (Santa Cruz Biotechnology), anti-SARS CoV-2 nucleocapsid mouse monoclonal (1:10) (Thermo Fisher Scientific) and anti-ACE2 mouse monoclonal (1:20) (R&D Systems). The next day, the inserts were washed with the IF washing buffer and incubated with an anti-mouse AlexaFlour 488-conjugated Ab (1:200) (Thermo Fisher Scientific) and/or an anti-rabbit AlexaFlour 647 (1:200) (Thermo Fisher Scientific) in IF washing buffer for 3 hours in the dark at 4°C. The cells were washed twice with PBS and incubated with rhodamine phalloidin (1:500) (Cytoskeleton Inc.) in IF washing buffer for 30 minutes at room temperature in the dark. After two washes with PBS, the cell nuclei were stained with NucBlue Fixed Cell Stain ReadyProbes (Thermo Fisher Scientific) for 30 minutes in the dark at room temperature. The epithelium was mounted on Tech-Med microscope slides (Thomas Scientific) using ProLong-Gold anti-fade mounting medium (Thermo Fisher Scientific). Images were captured using a confocal laser-scanning microscope (Olympus FV3000) enabled with a 60X objective. The 405-nm laser was used to excite the DAPI signal for nucleus detection, the 488-nm laser was used to excite Alexa Flour 488 for MUC5AC, ACE2, TMPRSS2 or SARS CoV-2 nucleocapsid protein detection, the 561-nm laser was used to excite rhodamine phalloidin for F-actin detection, and the 640-nm laser was used to excite Alexa Flour 647 for MUC5B, acetyl-alpha-tubulin, E-cadherin, ZO-1 or MUC5AC detection. At least two random fields were selected per sample and imaged. The images were processed with Imaris software version 9.5.1 (Oxford Instruments Group) and used for the conversion of Z-stack images (.oir format) to -tiff format and for additional image postprocessing. Separately, the tiling of Z-stack (3×3) images was captured using a Leica DMi8 microscope followed by image processing using a 3D deconvolution image processing module in the LASX software (Leica Microsystem) associated with the microscope. We used nine independent images to quantify the total cell number based on F-actin (Texas Red channel) and goblet cells using the Alexa Flour 647 channel (anti-MUC5B) with the Fiji multipointer option. Similarly, we quantified the MUC5AC^+^ cells using at least two confocal images.

### Immunohistochemistry

Healthy and COPD airway epithelia (mock or infected) were sectioned into 5-μm sections as described previously for immunohistochemistry. Before staining, the antigen retrieval process was performed using R-buffer A in a Retriever 2100. The slides were allowed to cool in the buffer overnight and washed three times in PBST (0.05% Tween-20) for 10 minutes. Using a Pap pen, a hydrophobic barrier was drawn around the tissue. Once the hydrophobic barrier was dry, the tissue was incubated with blocking buffer (10% goat serum in PBS) for 2 hours in a humidified, light-protected chamber. After incubation with blocking buffer, the tissue was immediately incubated with the following primary antibodies overnight at 4°C in a humidified, light-protected chamber: SARS-CoV-2 N (1:50) (Thermo Fischer Scientific) for the detection of SARS-CoV-2 nucleoprotein (N), SARS-CoV-2 spike (S) protein (1:100) (Thermo Fischer Scientific) for the detection of SARS-CoV2 S protein, acetylated-alpha-tubulin (1:500) (Cell Signaling Technologies) for the staining of ciliated cells, MUC5AC (1:500) (Cell Signaling Technologies) and MUC5B (1:500) (Atlas Antibodies) for the staining of goblet cells, and P63 (1:100) (Abcam Inc.) for the staining of basal cells. The next day, the slides were washed three times with PBST and then incubated for 45 minutes with anti-mouse Alexa Fluor 488 (Thermo Fisher Scientific) and anti-rabbit Alexa Fluor 647 (Thermo Fisher Scientific) secondary antibodies in a humidified, light-protected chamber. The slides were subsequently washed three times with PBST, incubated with rhodamine phalloidin (1:100) (Cytoskeleton Inc.) for 30 minutes at room temperature in a humidified, light-protected chamber, and washed three times with PBST. The nuclei were then stained by incubation with NucBlue Fixed Cell Stain ReadyProbes (Thermo Fisher Scientific) for 5 minutes at room temperature in a humidified, light-protected chamber. After incubation with the nuclear dye, the slides were washed once with PBST and twice with DI water, and coverglass was then placed on the tissue section using ProLong-Gold anti-fade mounting medium (Thermo Fisher Scientific). The slides were scanned using a Leica DMi8 inverted fluorescence microscope with a 63X oil objective, and the images were further processed using a 3D deconvolution image processing module in the LASX software (Leica Microsystem) associated with the microscope.

### Ciliary beat frequency (CBF)

Cilia on the apical surface of the cells in the differentiated epithelial layer (after 4 weeks of differentiation) were visualized in the phase-contrast mode with a Leica DMi8 microscope with a 20X objective and an attached environment control chamber (37°C with 5% CO_2_) (Leica Microsystems). For each Transwell, six different random fields were recorded for approximately 2.1 seconds at 120 frames per second. The images were captured at 37°C and analyzed using the Sisson-Ammons Video Analysis (SAVA) system V.2.1.15 to determine the CBF (Hz) (Ammons Engineering).

### Transepithelial electrical resistance (TEER)

The permeability of the differentiated epithelial layer (after 4 weeks of differentiation) was determined by measuring the TEER using an epithelial volt-ohm meter (EVOM2, World Precision Instruments, Inc.). The EVOM2 was calibrated according to the manufacturer’s instructions, and the STX2 electrode was sterilized with 70% ethanol before use. The internal electrode (smaller in size) was placed in the apical part of each Transwell (PBS was added during the TEER reading), and the external electrode (larger in size) was placed in the basal part of the Transwell, which contained ALI basal medium, to measure the membrane voltage and resistance of the epithelial layer. An empty Transwell insert (filled with PBS) containing no NHBE cells was used to correct for the background resistance. Three readings were taken for each Transwell. The TEER value of each sample was calculated by subtracting the background value.

### Quantitative real-time PCR

Airway epithelia cultured on 6.5-mm Transwell membranes were washed and treated for 1 minute at RT with RLT buffer (Qiagen) with 1% β-mercaptoethanol (Sigma-Aldrich). The cells were scraped using a cell scraper, collected into a QIAshredder tube and centrifuged at 15,000 rpm and 4°C for 3 minutes. The eluate was used for the extraction of total RNA using a Total DNA/RNA Extraction Kit (Qiagen), and DNase I treatment was performed to remove DNA from the samples according to the manufacturer’s instructions. We also followed a similar approach for the extraction of RNA from A549 cells. The RNA concentration was determined with an Epoch microplate spectrophotometer (BioTek). Five hundred nanograms of RNA was used for first-strand cDNA synthesis (Thermo Fisher Scientific) using Oligo(dT) primers (Thermo Fisher Scientific). qRT-PCR was performed using TaqMan assays (ACE2: Hs1085333_m1 and ACTB: Hs99999903_m1, for calibration) (Thermo Fisher Scientific) with the CFX384 Real-Time PCR System (Bio-Rad), and fold changes were calculated to determine the relative expression levels.

### Western blotting

The airway epithelium cultured on 6.5-mm Transwell membranes was washed with PBS, scraped out of all the cells, and pelleted by centrifugation at 10,000 rpm for 5 minutes. The cell pellet was incubated with 1x LDS loading buffer (Thermo Fisher Scientific) with proteinase inhibitor (Roche), transferred into a QIAshredder microcentrifuge tube, and centrifuged for 3 minutes at 15,000 rpm in a tabletop centrifuge. The elusion from the QIAshredder was collected and stored in a freezer. The protein concentration was measured using a BCA protein assay kit (Thermo Fisher Scientific). For the detection of ACE2, total protein (30 μg) was separated on 4-12% Bis-tris SDS polyacrylamide gels (reducing) and then subjected to dry blot transfer onto PVDF membranes according to the manufacturer’s instructions (Life Technologies). The PVDF membrane was imaged using an Odyssey CLX system (Li-Cor Biosciences). ACE2 was detected by Western blotting using anti-ACE2 goat polyclonal antibody (R&D Systems) and corresponding donkey anti-goat IRDye 800 secondary antibodies (Li-Cor Biosciences). For the loading control, alpha-tubulin was detected by anti-alpha-tubulin mouse monoclonal antibody (Sigma-Aldrich) and the corresponding goat anti-mouse IRDye 680 secondary antibody (Li-Cor Biosciences). Image Studio 5.2 (LI-COR Biosciences) was used to quantify the protein signal.

### Epithelial height measurement

Microscopic images of the IHC (described above) were used to quantify epithelial height by using the scale feature in LASX software of Leica DMi8 microscope. We used at least 4 independent slides for the measurement, at least three independent reads per slide. The plastic membrane of the Transwell was not included in the height measurement.

## QUANTIFICATION AND STATISTICAL ANALSYSIS

### Statistical analysis

Parameters such as the number of replicates, number of independent experiments, mean +/-SEM, and statistical significance are reported in the figures and figure legends. A p-value less than 0.05 was considered to indicate significance. Where appropriate, the statistical tests and post hoc statistical analysis methods are noted in the figure legends or Methods section.

**Figure S1.**
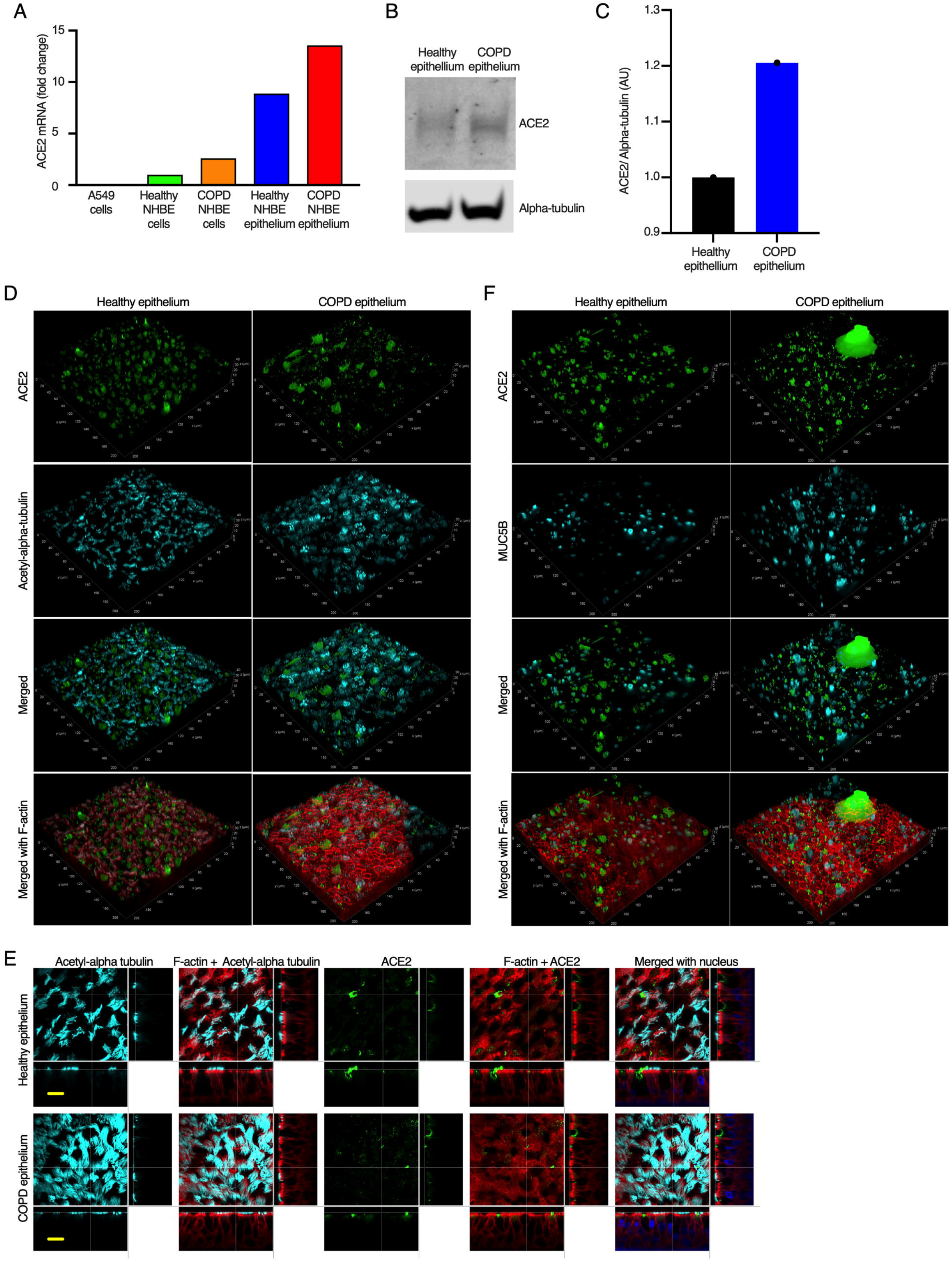
ACE2 expression is higher in the COPD airway epithelium. **(A)** ACE2 mRNA expression in total RNA extracted from the lung epithelial cell line (A549 cells) and primary NHBE cells from healthy adults and patients with COPD in monolayers or differentiated into airway epithelia was quantified by real-time PCR. The data were plotted as expression levels normalized to those in the NHBE healthy monolayer. The data were obtained by combining the quadruplet technical replicates of each sample. The graph represents the results from two independent real-time PCR runs. **(B).** ACE2 was detected in the airway epithelium (obtained after 4 weeks of differentiation of NHBE cells) by Western blotting. **(C).** The ACE2 signal (shown in B) was quantified (normalized to the alpha-tubulin level) and plotted relative to that in the healthy epithelium. **(D).** The apical sites of the airway epithelia were fixed and stained for ACE2 (anti-ACE2, green), cilia (anti-acetyl-alpha tubulin, cyan) and F-actin (rhodamine phalloidin, red). Deconvoluted Z-stack images are presented in the 3D view. **(E).** The apical sites of the airway epithelia were fixed and stained for cilia (anti-acetyl-alpha tubulin, cyan), ACE2 (anti-ACE2, green), F-actin (rhodamine phalloidin, red) and nucleus (DAPI, blue). Bar = 15 μm. **(F).** The airway epithelia (described in D) were stained for ACE2 (anti-ACE2, green), MUC5B (anti-MUC5B, cyan) and F-actin (rhodamine phalloidin, red). Deconvoluted Z-stack images are presented in the 3D view.

**Figure S2.**
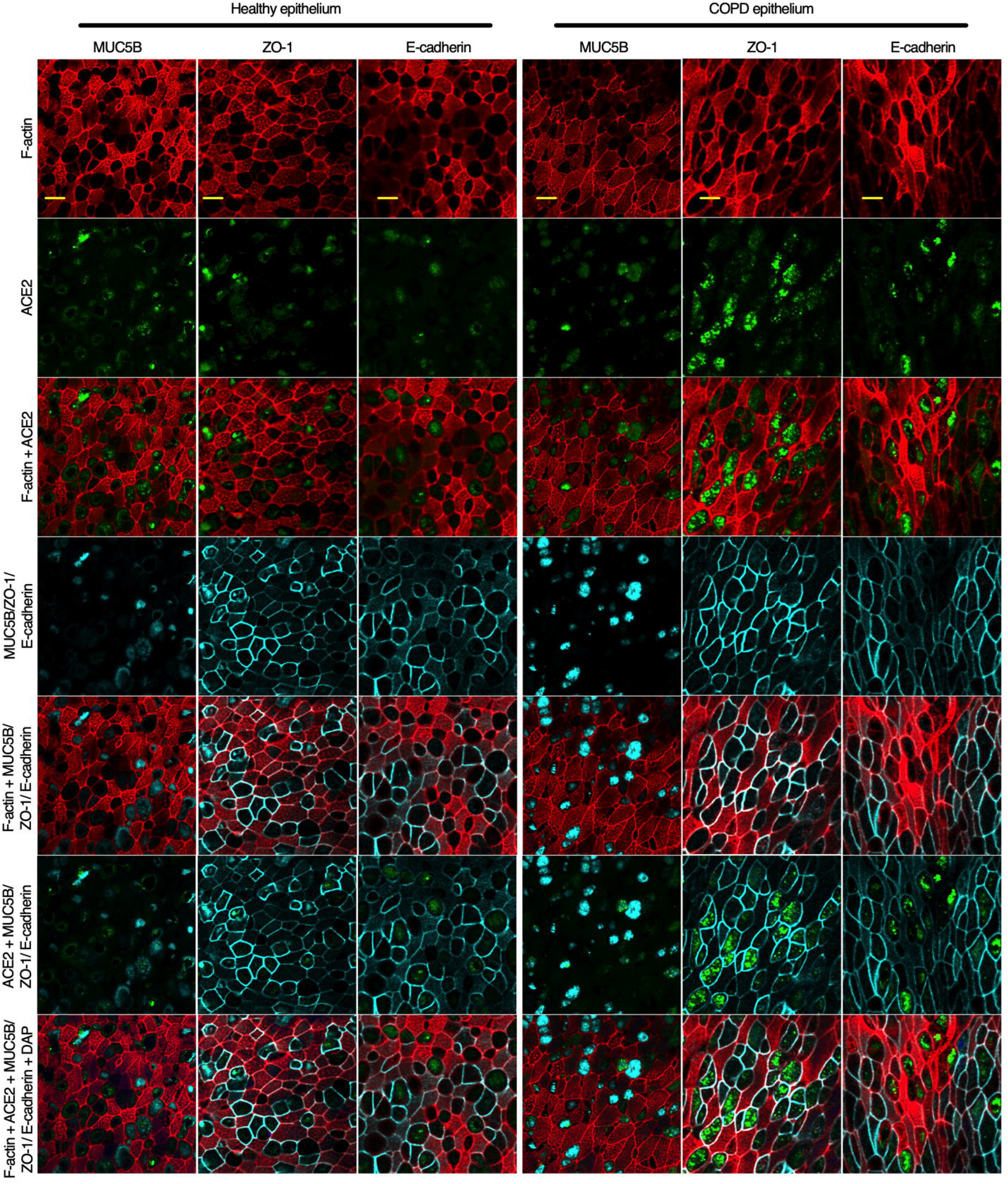
ACE2 expression within the goblet cell boundary. The airway epithelia (obtained after 4 weeks of differentiation of NHBE cells) were fixed and stained for ACE (anti-ACE2 antibody, green) and MUC5B (anti-MUC5B, cyan) or ZO-1 (anti-ZO-1 antibody, cyan) or e-cadherin (anti-e-cadherin antibody, cyan). F-actin (red) and nuclei (blue) were stained with rhodamine phalloidin and DAPI, respectively. The images represent multiple random areas obtained from an independent experiment. Bar = 10 μm.

**Figure S3.**
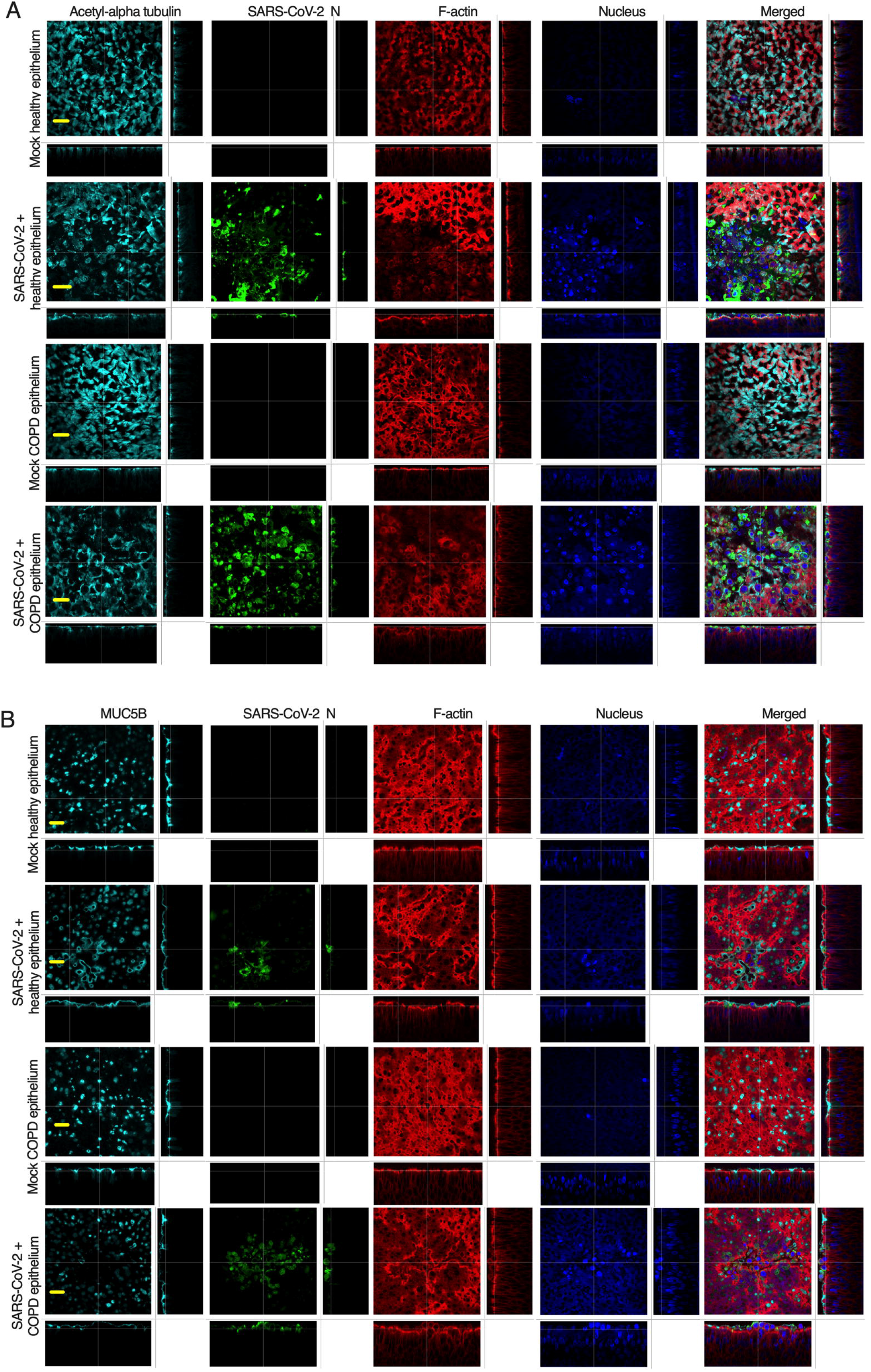
SARS-CoV-2 induces a cytopathic effect in the airway epithelia. The airway epithelia were mock-infected or infected with SARS-CoV-2 at an MOI of 0.1. At 4 days post infection (DPI), the epithelia were fixed, embedded in paraffin, sectioned, and stained for the basal cell marker P63 (anti-P63, green), SARS-CoV-2 spike (S) (anti-S, cyan), and F-actin (rhodamine phalloidin, red); the nuclei were stained with DAPI (blue). Bar = 20 μm.

**Figure S4.**
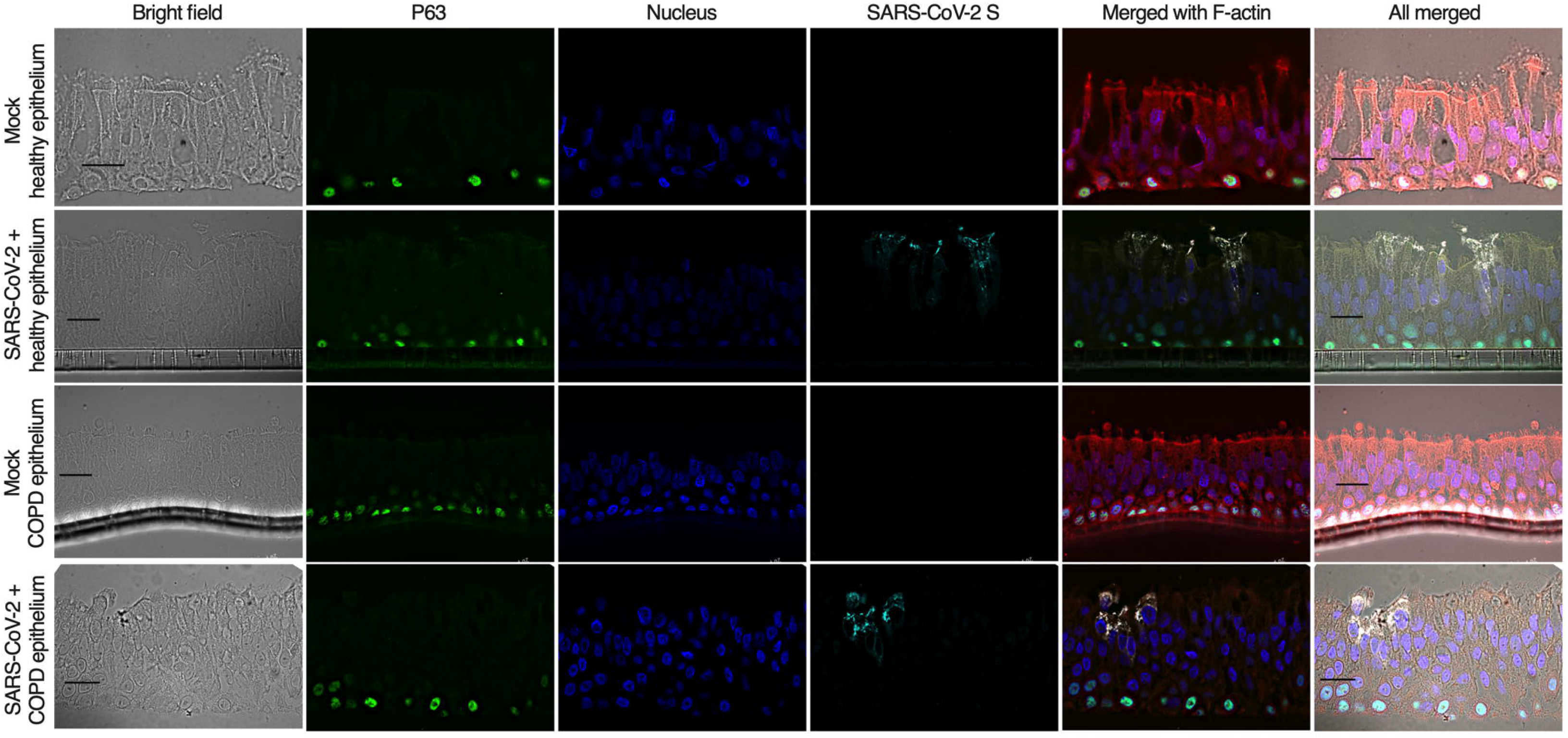
SARS-CoV-2 does not infect basal cells in the airway epithelia. The airway epithelia were mock-infected or infected with SARS-CoV-2 at an MOI of 0.1. At 4 days post infection (DPI), the epithelia were fixed, permeabilized, and stained for cilia (anti-acetyl-alpha tubulin, cyan), SARS-CoV-2 N (anti-N, green), and F-actin (rhodamine phalloidin, red); the nuclei were stained with DAPI (blue). The images represent multiple independent random areas from an independent experiment. Bar = 30 μm.

**Figure S5.**
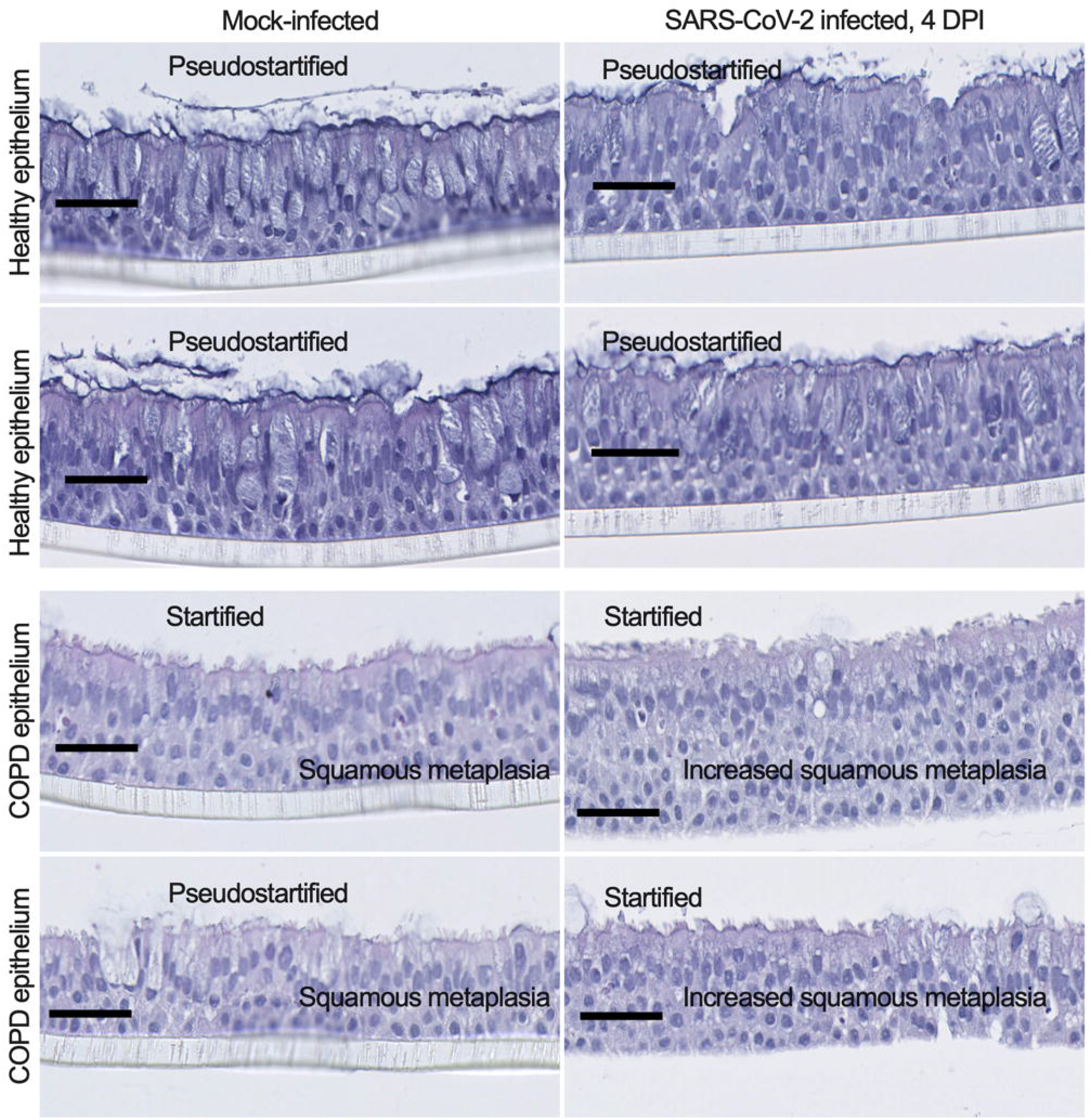
SARS-CoV-2 increases squamous metaplasia in the COPD airway epithelia. The airway epithelia were mock-infected or infected with SARS-CoV-2 at an MOI of 0.1. At 4 days post infection (DPI), the epithelia were fixed, permeabilized, and stained with hematoxylin and eosin (H&E). Bar = 50 μm.

---

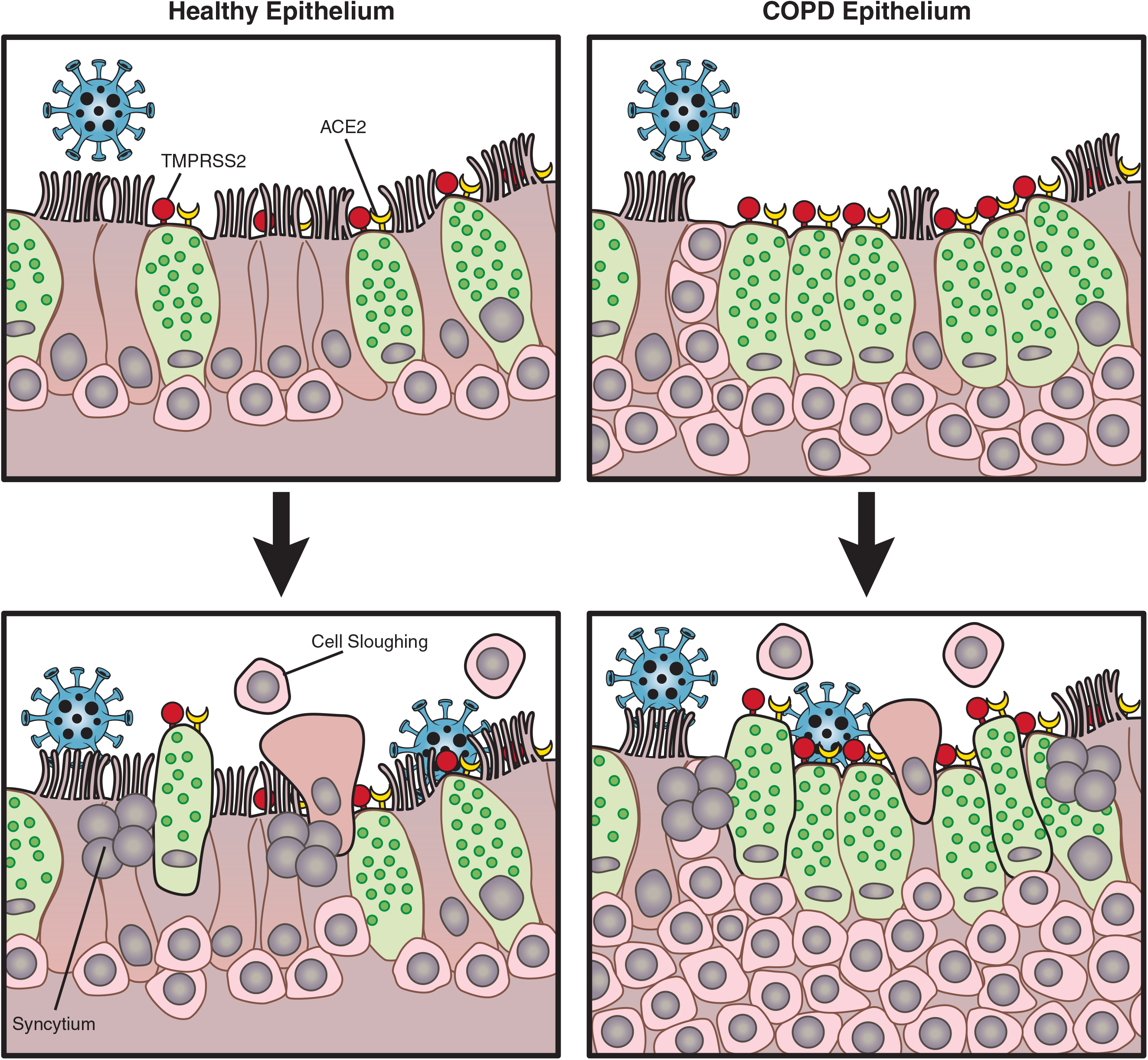

